# Tissue-specific genomic and transcriptomic dynamics of the OsHV-1 lytic cycle in the Pacific oyster *Magallana gigas* revealed by long-read sequencing

**DOI:** 10.64898/2026.11.15.766776

**Authors:** Aurélie Dotto-Maurel, Benjamin Morga, Isabelle Arzul, Germain Chevignon

**Affiliations:** Ifremer, ASIM, Adaptation Santé des Invertébrés Marins, F-17390 La Tremblade, France

**Keywords:** OsHV-1, *Magallana gigas*, herpesvirus, lytic cycle, long-read sequencing, transcriptome, tissue tropism

## Abstract

Ostreid herpesvirus 1 (OsHV-1) causes recurrent mass mortality of juvenile Pacific oysters (*Magallana gigas*) worldwide, yet its lytic cycle remains poorly characterized *in vivo*. In the absence of a permissive cell line, most studies have relied on intramuscular injection, which bypasses the natural route of infection, or on primary haemocyte cultures, which cannot capture tissue-level host-virus interactions. In addition, gene expression studies have relied on short-read sequencing, which fragments transcription units and cannot resolve complex transcriptional architectures. Long-read sequencing has recently revealed abundant overlapping, antisense and polycistronic transcripts in other herpesviruses, but whether OsHV-1 shares this complexity, and how its transcriptional program varies across host tissues during natural infection, remains unknown. Here, we exposed naive oysters to OsHV-1 by bath exposure and combined long-read genomic and transcriptomic sequencing of gills, mantle, digestive gland and adductor muscle at increasing levels of infection. OsHV-1 genomes were assembled directly from infected tissues, and intra-host viral diversity was structured by host rather than tissue. We identified 410 unspliced viral transcripts, including overlapping, antisense and polycistronic transcripts. The viral transcriptional program was largely conserved across tissues and resembled that previously described in haemocytes in vitro. In the host, infection triggered a response shared by all tissues, but earlier and stronger in the gills and mantle. Together, these results provide the first long-read view of the OsHV-1 lytic cycle in its natural host.

**IMPORTANCE:** OsHV-1 is one of the main pathogens threatening Pacific oyster aquaculture. However, the lack of permissive cell lines has limited our understanding of how it replicates in its host. By infecting oysters via seawater and sequencing the full-length genomes and transcripts of the virus directly from infected tissues, we demonstrate that OsHV-1 has a complex transcriptional architecture, similar to human herpesviruses. Furthermore, we show that its replication program is remarkably uniform across tissues. We also identify a small set of poorly characterized viral genes that are highly expressed early in infection and represent promising targets for functional studies. These data provide a foundation for improving OsHV-1 genome annotation and understanding herpesvirus biology beyond vertebrates.

## INTRODUCTION

Since the 1990s, the Pacific oyster (*Magallana gigas*, formerly *Crassostrea gigas*), has experienced recurrent outbreaks caused by the herpesvirus *Ostreavirus ostreidmalaco1* (OsHV-1) (Davison et al., 2005; Nicolas et al., 1992; Renault et al., 1994, 2001). Howewer, OsHV-1 and its variants have also been detected in several other marine mollusc species, including the scallop *Azumapecten farreri* (Bai et al., 2019), the ark clam *Anadara broughtonii* (Xia et al., 2015), and the flat oyster *Ostrea edulis* (Arzul, Renault, et al., 2001; Pelletier et al., 2025; Renault et al., 2000). In addition, herpes-like viruses resembling OsHV-1 but not characterized phylogenetically have been reported in the clams *Ruditapes philippinarum* and *Ruditapes decussatus* (Renault, 1998; Renault et al., 2001; Xia et al., 2015), the scallop *Pecten maximus* (Arzul, Nicolas, et al., 2001), and the Portuguese oyster *Magallana angulata* (Arzul, Renault, et al., 2001). To date, OsHV-1 has been reported across various oyster-farming regions worldwide, from Europe to Asia, Oceania and the Americas (Abbadi et al., 2018; Barbieri et al., 2019; Batista et al., 2015; Burge et al., 2006; Burioli et al., 2017; Clegg et al., 2014; Fabioux et al., 2002; Gittenberger et al., 2016; Grijalva-Chon et al., 2013; Hine & Thorne, 1997; Hwang et al., 2013; Mello et al., 2018; Meyers et al., 2009; Mortensen et al., 2016; Moss et al., 2007; Webb et al., 2007).

Like other herpesviruses, OsHV-1 life cycle is thought to alternate between lytic and persistent phases. Some authors have shown that mortality of juvenile oysters is observed once seawater temperature exceeds 16°C suggesting that the virus lytic phase is activated through this temperature increase (Dégremont, 2013; Petton et al., 2013; Renault et al., 1994, 2011; Segarra et al., 2014). Conversely, when temperatures fall below approximately 13°C, no oyster mortality is recorded and OsHV-1 generally no longer detected by quantitative PCR (qPCR), suggesting that the virus shifts to a persistent phase (Dégremont et al., 2013; Degremont & Benabdelmouna, 2014; Evans et al., 2017; Petton et al., 2013).

To date, the mechanisms regulating the OsHV-1 lytic phase in the Pacific oyster are not fully understood. The virus is thought to infect oysters through pallial organs, particularly *via* the gills, before reaching the haemolymphatic system, which would allow the virus to spread to various oyster tissues (Martenot et al., 2016; Schikorski, Faury, et al., 2011). Transmission electron microscopy has identified viral particles in fibroblast-like cells in several connective tissues of infected larvae and spat, including the mantle, labial palps, gills and digestive gland, demonstrating that OsHV-1 could replicate in these tissues (Nicolas et al., 1992; Renault et al., 1994, 2001). These findings are consistent with results obtained by *in situ* hybridization and immunohistochemistry, detecting OsHV-1 DNA, RNA and proteins not only in these organs but also in the adductor muscle, heart, gonad and neural ganglia (Arzul et al., 2002; Corbeil et al., 2015; Lipart & Renault, 2002; Martenot et al., 2016; Schikorski, Faury, et al., 2011). Furthermore, reverse transcription quantitative PCR (RT-qPCR) has revealed that OsHV-1 gene expression can be detected very early in the infection process, from two hours post-infection (hpi) in mantle tissue and as early as one hpi in primary haemocyte cultures (Morga et al., 2017; Segarra et al., 2014).

Recently, short-read RNA-seq has substantially advanced our understanding of OsHV-1 gene expression during the lytic phase. In particular, a dual RNA-seq analysis of haemocytes infected *in vitro* revealed a temporally regulated viral transcriptional cascade, consistent with the expression patterns observed in other herpesviruses, and identified five genes expressed at exceptionally high levels during the first 24 h of the lytic cycle (Dotto-Maurel, Luyer, et al., 2025). A similar approach applied to an OsHV-1 genotype infecting ark clams (*Scapharca broughtonii*) also identified several highly expressed genes, although their temporal dynamics were not investigated (Bai et al., 2018). Short-read technologies, however, fragment transcription units and are therefore poorly suited to resolving the often complex architecture of viral transcription units. Long-read sequencing has emerged as a powerful alternative, capturing full-length RNA molecules and thereby providing a more accurate and comprehensive view of herpesvirus transcriptome architecture (Kakuk et al., 2021; Tombácz et al., 2016, 2018, 2019). Applied to several herpesviruses, this approach has revealed transcriptomes far more complex than previously thought, including overlapping transcriptional units, multigenic transcripts and antisense transcripts (Boldogkői, Moldován, et al., 2019). In OsHV-1, long-read transcriptomic analysis of the clam-infecting genotype has likewise revealed an additional layer of complexity, including polycistronic and antisense transcripts. (Bai et al., 2021; Rosani et al., 2024). However, the temporal dynamics of these transcripts throughout the lytic cycle remain unclear, and this genotype does not infect *M. gigas*, for which long-read transcriptomic data is still lacking.

From a genomic perspective, long-read sequencing is equally well suited to resolving the complex genome structure of herpesviruses such as OsHV-1 (Dotto-Maurel et al., 2022; Dotto-Maurel, Pelletier, et al., 2025). The OsHV-1 genome displays the canonical herpesvirus architecture, with two unique regions, U_L_ (unique long, 170 kb) and U_S_ (unique short, 3.5 kb), each flanked by inverted repeats (TR_L_/IR_L_, 7.5 kb each, and TR_S_/IR_S_, 9.7 kb each) and separated by a third unique region, X (1.5 kb), giving the overall organization TR_L_-U_L_-IR_L_-X-IR_S_-U_S_-TR_S_ (Davison et al., 2005). As in other herpesviruses, this genome does not exist as a single molecular form. Four major isomers coexist at equimolar levels within the same sample, differing in the relative orientation of U_L_ and U_S_ (P: prototype; I_L_: U_L_ inverted; I_S_: U_S_ inverted; I_LS_: both inverted), and each of them occurs in three additional minor forms defined by the position of the X region, which may be duplicated (D_X_), translocated to the left genome terminus (T_X_) or absent (N_X_) (Dotto-Maurel et al., 2022; Dotto-Maurel, Pelletier, et al., 2025). Long-read sequencing now allows these configurations to be characterized directly, yet whether this structural plasticity has any functional relevance during the lytic phase remains an open question (Dotto-Maurel, Arzul, et al., 2025).

A further obstacle to the study of OsHV-1 is the absence of a permissive cell line, which leaves two imperfect experimental alternatives. *In vivo*, most studies of the lytic phase have relied on oysters infected by intramuscular injection (Petton et al., 2013; Renault et al., 1994, 2011; Segarra et al., 2014). This route offers a high level of experimental control but bypasses the natural infection process, in which the virus is transmitted between oysters, particularly larvae and juveniles, through cohabitation with infected individuals or exposure to virus-contaminated seawater (Schikorski, Faury, et al., 2011). *In vitro*, primary haemocytes cultures have provided valuable insight into replication dynamics (Dotto-Maurel, Le Luyer, et al., 2025; Morga et al., 2017), but they cannot reproduce the tissue-level host-pathogen interactions that shape infection in the whole animal.

Here we combined a balneation-based infection protocol, which reproduces the natural route of exposure of the Pacific oyster *M. gigas* to OsHV-1, with sampling every 3 h over a 99-h period and multi-omics Oxford Nanopore long-read sequencing. Four tissues, in which infection is well documented, were analyzed: gills, mantle, digestive gland and adductor muscle (Lipart & Renault, 2002; Martenot et al., 2016). This design allowed us to resolve the transcriptomic dynamics of the lytic cycle at full-length resolution, to track the plasticity of the viral genome architecture over the course of infection, and to assess tissue-specific host responses.

## RESULTS

### 1. OsHV-1 infection showed a consistent temporal progression

In the infected control tank, mortality began at 78 hours post-infection (hpi), and cumulative mortality reached 64% at the end of the experiment (Figure 1.A). In the three experimental tanks, mortality started slightly later, between 81 and 84 hpi. Moribund oysters carried more than 10⁵ OsHV-1 genome copies per ng of DNA (data not shown), consistent with active viral replication. In the non-infected control tank, no mortality nor detection of OsHV-1 DNA was observed (Figure 1.A).

**Figure 1:**
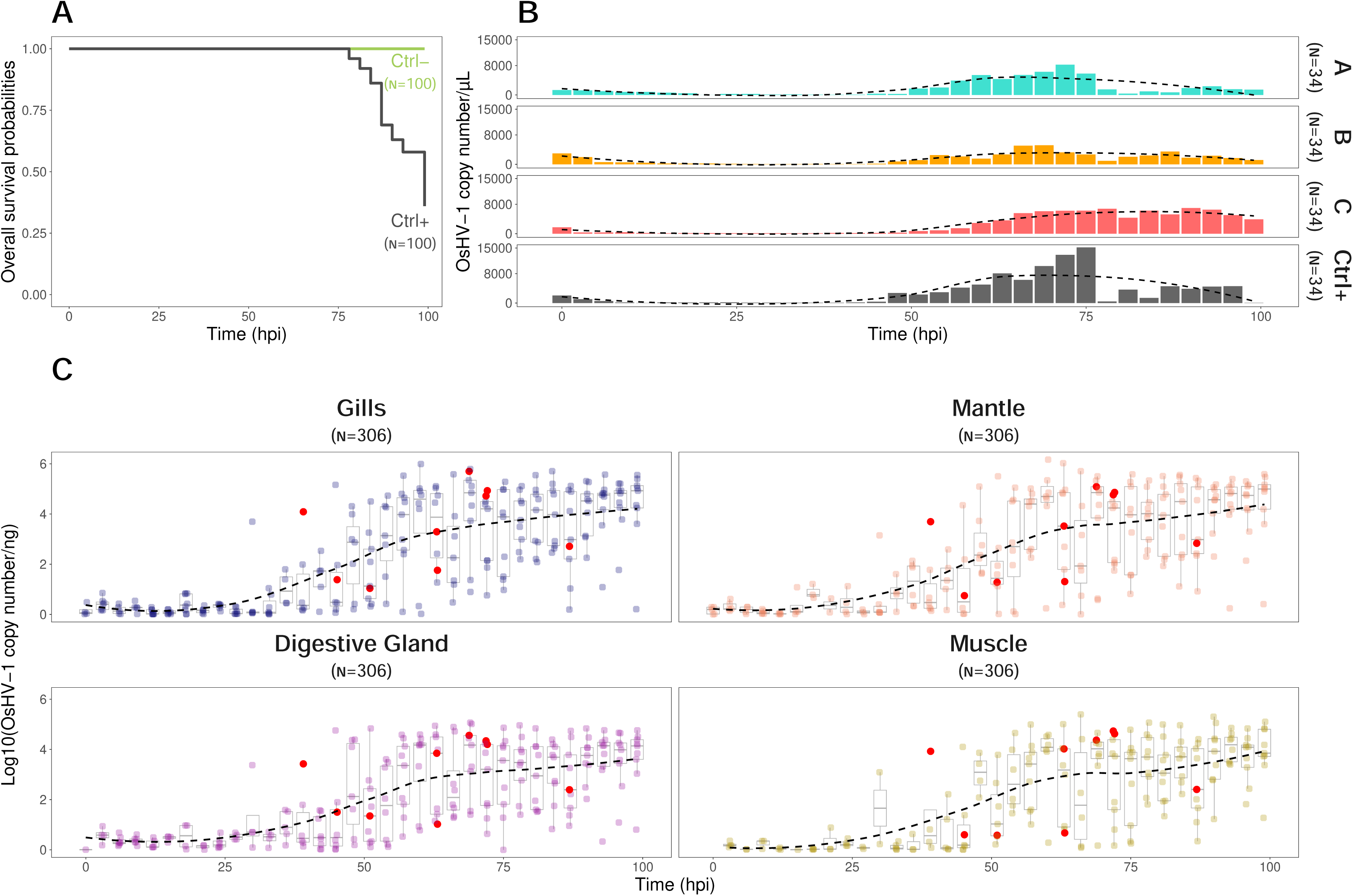
OsHV-1 infection dynamics and oyster survival over time. A. Kaplan-Meier curve showing the overall survival probability of infected (Ctrl +) and not infected control tank (Ctrl-) oysters over time in hours post-infection (hpi). B. Temporal dynamics of OsHV-1 DNA detected in seawater from experimental tanks (A, B, C) and infected control tank (Ctrl+). The Y-axis indicates the OsHV-1 copy number per µL and the X-axis shows time in hpi. The dotted black line indicates the smoothed trend fitted to the data. C. OsHV-1 DNA levels in the gills, mantle, digestive gland and muscle over time. The y-axis represents the log₁₀ of OsHV-1 copies per ng of DNA, and the x-axis indicates time in hpi. Each dot in the graph represents a replicate. The dotted black line indicates the smoothed trend fitted to the data. Red dots represent sequenced samples.

The dynamics of OsHV-1 DNA in seawater were similar in all exposed tanks (Figure 1.B). At 0 hpi, seawater contained a mean of 2 × 10³ OsHV-1 genome copies per µL of seawater. This concentration decreased sharply between 0 and 3 hpi and then remained very low or below the limit of detection until 51 hpi. OsHV-1 DNA concentration then increased again to 72 hpi, decreased at 78 hpi, and rose slightly towards the end of the experiment.

In oyster tissues, OsHV-1 DNA became detectable at around 24 hpi and increased thereafter until the end of the experiment, with comparable dynamics in the four tissues analyzed (Figure 1.C). Viral load was nonetheless characterized by substantial inter-individual variability throughout the experiment, as reflected by a subset of oysters harboring very low OsHV-1 genome copy numbers even at the latest sampling time points (Figure 1.C).

To investigate the OsHV-1 lytic phase at both the genomic and transcriptomic levels, gills, mantle, digestive gland and muscle of 12 individuals were selected to represent four infection levels, used as a proxy for infection progression, namely not infected, lightly infected (∼10 OsHV-1 genome copies/ng DNA), moderately infected (∼10³ genome copies/ng DNA), and highly infected (∼10⁵ genome copies/ng DNA) (Figure 1.C, red dots; selected individuals are indicated in bold in Table S1).

### 2. Plasticity of OsHV-1 genomic architecture during the lytic cycle

#### 2.1. Proportion of OsHV-1 gDNA reads increases with infection level

A total of 20 gDNA libraries were sequenced across five ONT runs (4 tissues × 1 replicate for each of the lightly and moderately infected levels, and 4 tissues × 3 replicates for the highly infected level), yielding between 0.2 and 3.8 Gb per library (Table S1). Because ONT read lengths are highly variable, sequencing output was quantified as numbers of aligned bases rather than numbers of reads.

In lightly infected oysters, 0.004-0.031% of sequenced bases aligned to the OsHV-1 genome and more than 98% to the *M. gigas* genome. In moderately infected oysters, these proportions were 0.7-2% for OsHV-1 and more than 97% for *M. gigas*. In highly infected oysters, they were 0.6-14% and 83-98%, respectively (Figure S1.A, Table S1). Moreover, a small proportion of bases mapped to both genomes simultaneously (less than 0.00003%, 0.06%, and 0.5% for lightly, moderately, and highly infected samples, respectively), and approximately 1% remained unaligned (Figure S1.A, Table S1). Taxonomic classification of unaligned reads revealed a predominance of bacterial sequences and a minor viral fraction, consisting almost exclusively of bacteriophages (Supplementary File 1, Figure S2.A, Table S2).

#### 2.2. OsHV-1 reads enable complete genome assembly across tissues

The read N50 of OsHV-1-aligned reads ranged from 1,980 bp (mantle of a lightly infected oyster) to 9,379 bp (gills of a moderately infected oyster) (Figure 2). Mean N50 was 5,508 bp in the gills, 5,879 bp in the mantle, 5,214 bp in the digestive gland and 3,070 bp in the muscle.

**Figure 2:**
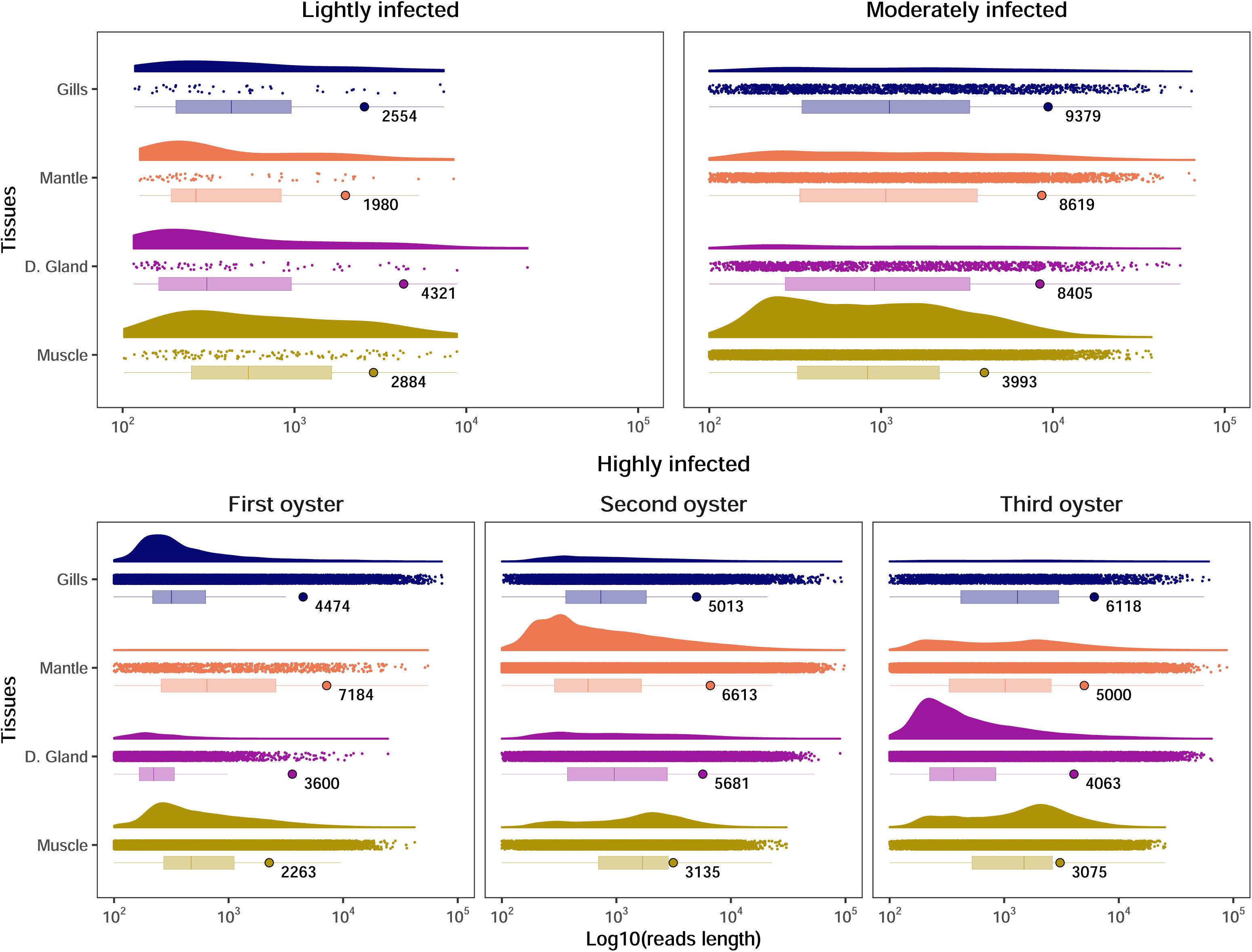
Raincloud plot showing the distribution of OsHV-1 read length across tissues and infection conditions. This figure shows gDNA reads that have been aligned to the OsHV-1 genome. The y-axis represents the different oyster tissues, and the x-axis shows the log₁₀-transformed OsHV-1 read length. For each tissue, the top layer shows the density distribution of read lengths, followed by a jitter plot of individual read lengths in the middle, and a boxplot at the bottom, with the N50 value indicated by a colored point and its corresponding value. Each panel corresponds to a different infection condition, ranging from low (left) to high (right). Note that three biological replicates are shown for the highly infected condition, as all three were sequenced to enable intra-condition variant calling (see Methods), whereas only one replicate was sequenced for the lightly and moderately infected conditions.

*De novo* assembly of the pooled reads not aligned to the *M. gigas* genome, from all tissues and infection levels, yielded two OsHV-1 contigs. Neither displayed the canonical OsHV-1 genome arrangement TR_L_-U_L_-IR_L_-X-IR_S_-U_S_-TR_S_ (Figure S3.A). The first contig (52,042 bp) was organized as TR_L_ - X - IR_S_ - U_S_ - IR_S_ - X - IR_L_ - U_L_, with a partial U_L_. The second (227,338 bp) was organized as TR_S_- U_S_ - IR_S_ - X - IR_L_ - U_L_ - IR_L_ - X - IR_S_ - U_S_ - IR_S_. The two contigs were merged into a single sequence with the canonical arrangement using a custom script. After polishing with Medaka, this yielded a complete OsHV-1 genome of 203,377 bp (Figure S3.A).

In a whole-genome phylogeny, based on non-redundant genomes, including the new genome and all publicly available OsHV-1 genomes, the new genome was most closely related to France_2021_OZ210309.1 (GenBank: GCA_964656485.1) and France_2020_OZ210743.1 (GenBank: GCA_964656475.1). Those genome were assembled from oysters collected in the same geographical area in 2021 and 2020 (Dotto-Maurel, Pelletier, et al., 2025) (Figure S3.B).

To validate the assembly, reads not aligned to *M. gigas* from each tissue and infection level were realigned to the new genome (Figure S3.C). In lightly infected oysters, the genome was only partially covered (20–45% of its length), with a mean depth of about 3× in the gills, mantle and digestive gland and 7× in the muscle. In moderately and highly infected oysters, the genome was fully covered. Mean depths ranged from 55× (gills) to 329× (muscle) in moderately infected oysters and from 274× (gills) to 983× (mantle) in highly infected oysters.

#### 2.3. Isomers-supporting sequences were identified in all tissues

The OsHV-1 genome exists as four major isomers, prototype (P), inverted L (I_L_), inverted S (I_S_), and inverted L and S (I_LS_). They differ in the relative orientations of U_L_ and U_S_ (Davison et al., 2005). A read can be assigned to one isomer only if it spans the internal repeat region IR_L_-X-IR_S_ and is anchored in both U_L_ and U_S_. OsHV-1-aligned reads longer than 9 kb were first extracted as candidates. Only one was obtained from lightly infected oysters (digestive gland), which were therefore excluded from this analysis. Moderately and highly infected oysters yielded 60 to 450 and 1,500 to 8,000 candidate reads per tissue, respectively.

In highly infected oysters, reads supporting each of the four major isomers were found in the gills (11 to 19 reads per isomer), the mantle (29 to 42) and the digestive gland (4 to 8) (Figure 3). No supporting read was found in the muscle, consistent with the shorter reads obtained from this tissue. In moderately infected oysters, supporting reads were scarce (1 to 8 per tissue), and the four isomers were therefore not all detected in each tissue.

**Figure 3:**
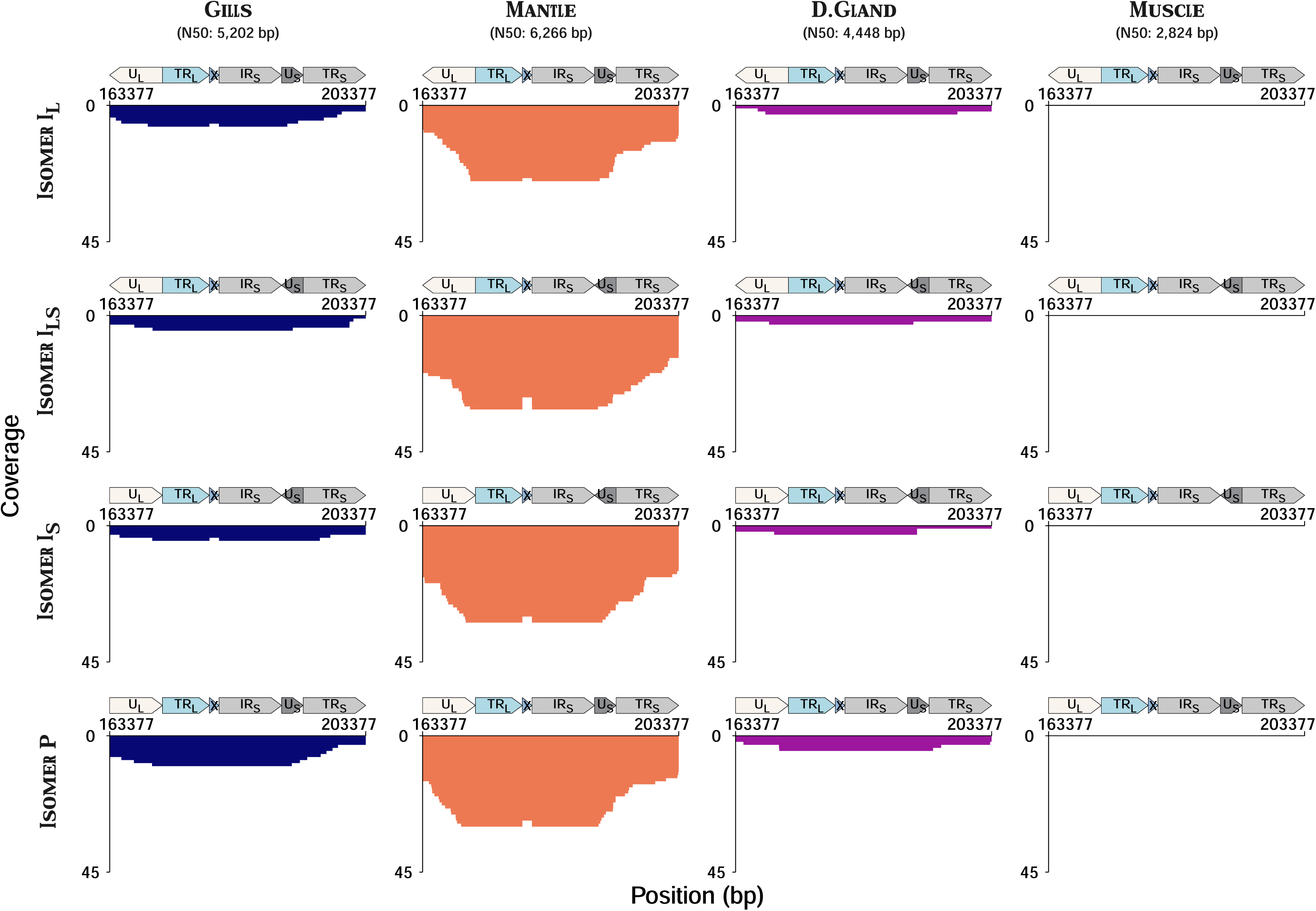
Coverage of OsHV-1 major isomers across tissues in the highly infected condition. Each column represents one of the four tissues and each row corresponds to one of the four major isomers. The y-axis shows the combined read coverage across the three biological replicates, while the x-axis represents the position along each isomer sequence in base pairs (bp).

In addition, 10 reads from the gills and mantle of highly infected oysters joined IR_L_ directly to IR_S_, without the intervening X region, consistent with the minor N_X_ isomer (X absent) or T_X_ isomer (X translocated), both of which produce this junction (Figure 3).

#### 2.4. OsHV-1 single nucleotide variants are not related to tissue types

Eleven variable positions were identified across the 12 libraries (Figure 4, Table S3). The number of variants ranged from 2 to 7 per oyster. Oyster n°1 carried 7 variants (VAF range 61.3-100%), of which 6 were detected only in this oyster and 4 were detected in all four of its tissues. Oyster n°2 carried 2 variants (VAF range 58.5-99.8%), both detected in all four tissues. One of these, at position 126340 bp, was also detected in the other two oysters. Oyster n°3 carried 4 variants (VAF range 65.9–88.6%), of which 3 were detected only in this oyster, and all 4 were detected in all four of its tissues (Figure 4, Table S3). Moreover, Jaccard dissimilarity averaged 0.07 ± 0.13 between tissues of the same oyster and 0.85 ± 0.04 between oysters for the same tissue. A PERMANOVA on Bray-Curtis dissimilarities of VAF profiles attributed 97.7% of the variance to oyster identity (R² = 0.977, F₂,₆ = 189.95, p = 0.001) and 0.8% to tissue (R² = 0.008, F₃,₆ = 1.04, p = 0.45), meaning that oyster identity, rather than tissue, accounted for almost all of the explained variance in viral variant composition.

**Figure 4:**
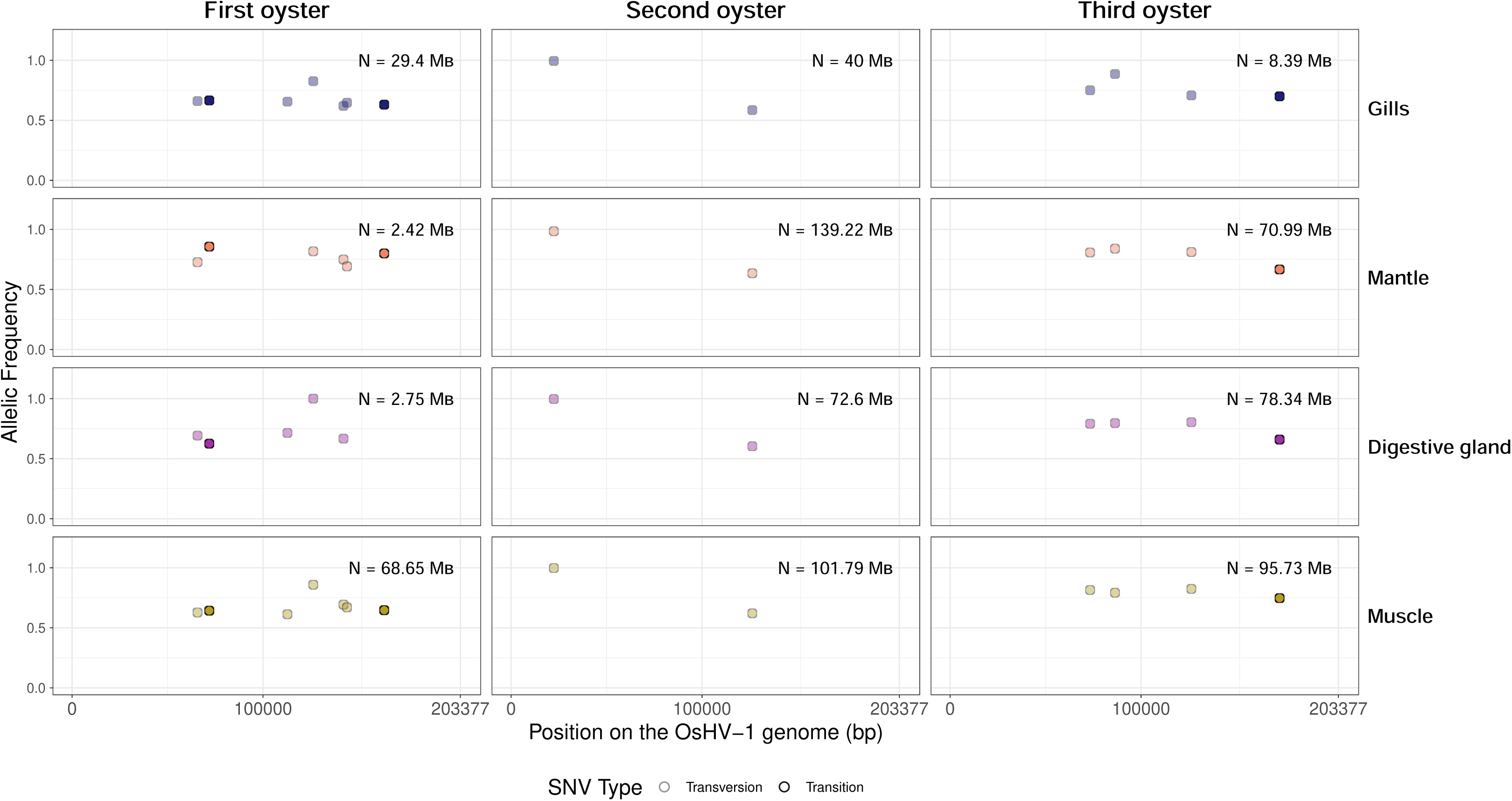
Distribution of SNVs across the OsHV-1 genome in the three highly infected oysters. Columns represent individual oysters, and rows correspond to the four sampled tissues (gills, mantle, digestive gland, and muscle). Each point represents a SNV, positioned according to its genomic location on the OsHV-1 reference genome (x-axis) and its allelic frequency (y-axis). Point fill indicates the SNV type, with transitions shown as opaque circles and transversions as semi-transparent circles. The total number of viral bases (N) used for SNV detection in each sample is indicated in the upper-right corner of each panel.

### 3. OsHV-1 gene expression dynamics during the lytic cycle

#### 3.1. OsHV-1 cDNA sequences are found in the four infection level for all tissues

A total of 48 cDNA libraries (4 infection levels × 4 tissues × 3 oysters) were sequenced across 23 ONT runs. Read yields per library ranged from 54,501 to 12,643,065 reads (0.027–9.3 Gb); the lowest-yield libraries corresponded to muscle samples, particularly in the not infected group (Table S1). No read aligned to the OsHV-1 genome in not infected oysters and approximately 99% of bases aligned to the *M. gigas* genome. In lightly infected oysters, 0.001-0.03% of bases aligned to OsHV-1 and approximately 99% to *M. gigas*. In moderately infected oysters, these proportions were 0.2-4% and 95-99%, respectively. In highly infected oysters, they were 0.5-20.5% and 79-99% (Figure S1.B, Table S1).

Reads aligning to both genomes accounted for at most 0.01% and 0.03% of bases in lightly, moderately and highly infected oysters, respectively. They were attributed to chimeras formed during library preparation. In each library, at most 1.55% of bases remained unaligned. As for gDNA libraries, taxonomic classification of these reads identified a small number of picorna-like RNA virus transcripts restricted to the digestive gland (Supplementary File 1, Figure S2.B, Table S2).

Read N50 ranged from 448 to 1,226 bp for OsHV-1-aligned reads (mean across libraries: 775 bp), from 244 to 879 bp for *M. gigas*-aligned reads (660 bp) and from 81 to 578 bp for unaligned reads (297 bp) (Table S1).

#### 3.2. Overall OsHV-1 expression profile is similar between tissues

cDNA reads from moderately and highly infected oysters were aligned to a non-redundant version of the new OsHV-1 genome. This version corresponds to the P isomer without its terminal repeats (U_L_-IR_L_-X-IR_S_-U_S_). OsHV-1 coverage profiles were similar across the four tissues, although coverage depth varied with tissue and infection level (Figure 5.A). Notably, ORF18, ORF107 (coding for proteins of unknown function with predicted intrinsically disordered regions), ORF27 (coding for putative dUTPase) and ORF80 and ORF120 (coding for encode putative membrane proteins) were the top 5 most highly expressed genes in both moderately and highly infected conditions across all tissues. Highest values were consistently observed in the mantle (Figure 5. A).

**Figure 5:**
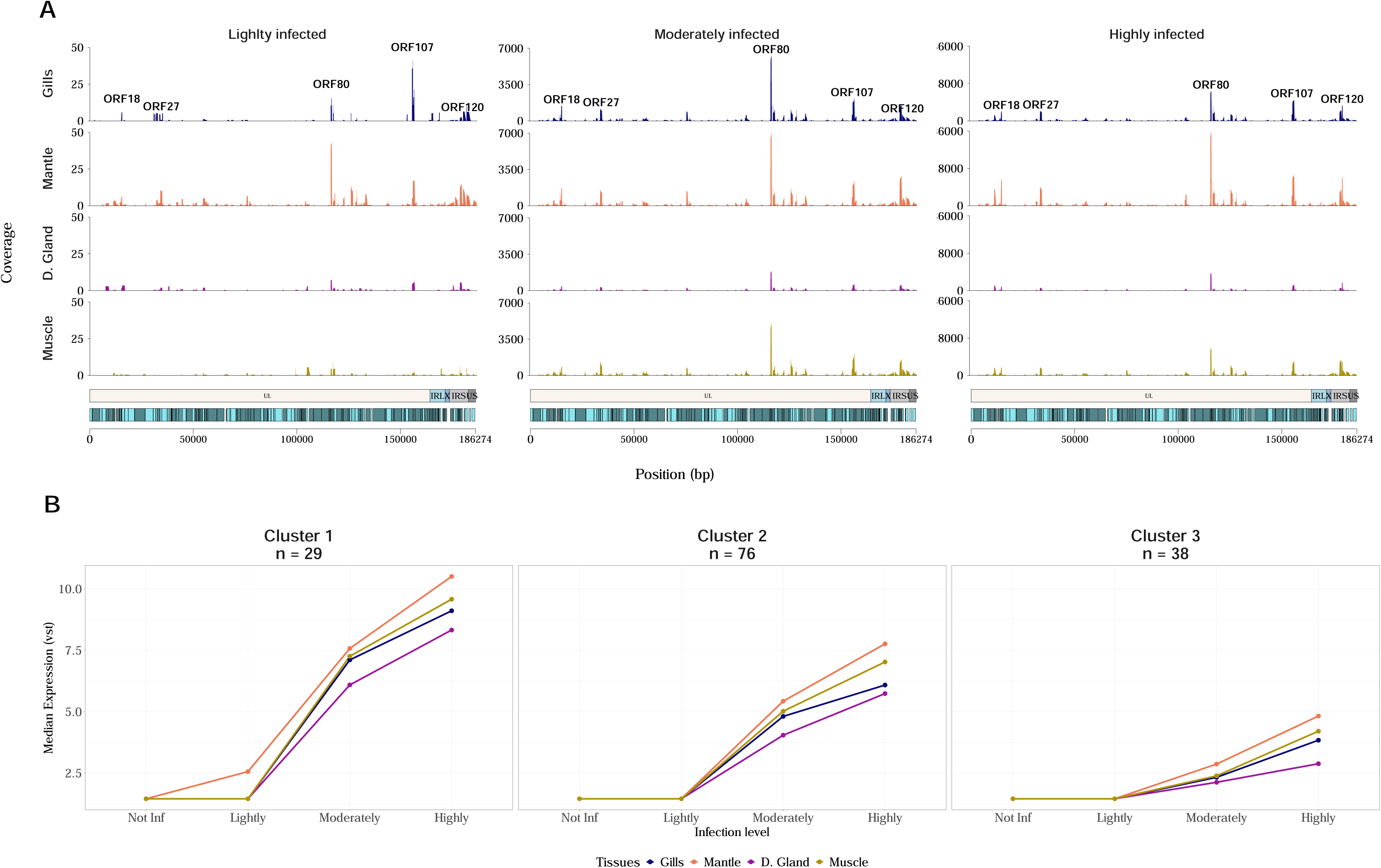
OsHV-1 genome-wide expression profiles across tissues, infection levels, and temporal gene-expression clusters. A. OsHV-1 whole genome expression across tissues and infection levels (n = 3 oysters per tissue and condition). The Y-axis shows the mean coverage per 100 bp windows for each infection level: lightly infected (left panel, 0x to 50x), moderately infected (middle panel, 0x to 7 000x) and highly infected (right panel, 0x to 35 400x). In each panel, the rows represent different tissues: gills (darkblue), mantle (orange), digestive gland (purple) and muscle (darkyellow). The X-axis shows the position of the OsHV-1 genome in base pairs. Above the X-axis, OsHV-1 genomic regions are annotated: U_L_ (white), IR_L_ (light blue), X (dark blue), IR_S_ (light grey) and U_S_ (dark grey). Open reading frames (ORFs) are also displayed: forward-directed ORFs are shown in dark blue and reverse-directed ORFs in light blue. B. Clustering of OsHV-1 genes based on their expression (n = 3 oysters per tissue and condition). Each panel shows one of the three clusters. The Y-axis shows the median expression, transformed using a variance-stabilizing transformation (vst), and the X-axis shows time in hours post-infection. Lines and dots are colored according to the tissues.

#### 3.3. OsHV-1 genes are expressed differently depending on infection level

143 OsHV-1 genes detected in at least one library across the dataset were clustered according to their median expression profile across infection levels, based on variance-stabilized values (Figure 5.B, Table S4). K-means clustering identified three groups differing in overall expression magnitude. Cluster 1 comprised 26 genes with the highest expression levels across all infection stages. It included the five most highly expressed ORFs as well as ORF87 coding for a putative inhibitor of apoptosis and ORF44 predicted to be involved in DNA replication. Cluster 2, the largest, comprised 76 genes with intermediate expression levels. It included ORF75, which code for putative dUTPase, and three genes coding for putative inhibitors of apoptosis (ORF42, ORF99 and ORF106). It also included seven of the nine genes predicted to be involved in DNA replication (ORF7, ORF20, ORF44, ORF49, ORF51, ORF66, ORF67 and ORF100). Cluster 3 comprised 38 genes, following a similar increasing trend as cluster 2 but reaching lower expression levels overall. It included ORF109, which codes for a putative DNA packaging protein.

### 4. Structure and expression of OsHV-1 transcripts

#### 4.1. Long-read sequencing identified 410 OsHV-1 transcripts

Aligning full-length cDNA reads to the non-redundant OsHV-1 genome identified 410 transcripts, ranging from 77 to 2,901 bp in length (median: 762 bp) (Figure 6.A, Table S4). Transcripts were classified according to the completeness and orientation of the ORF(s) they overlapped (Figure 6.B). Of these, 230 contained at least one incompletely covered ORF and were classified as partial, while the remainder were fully covered by one or more ORFs and classified as monocistronic (79), antisense (57), polycistronic (27), intergenic (9, no ORF overlap) or complex (8, ORFs in discordant orientations). Among partial transcripts, 171 contained a single incompletely covered ORF (partial monocistronic); the remaining subtypes are detailed in Figure 6.B and Table S4.

**Figure 6:**
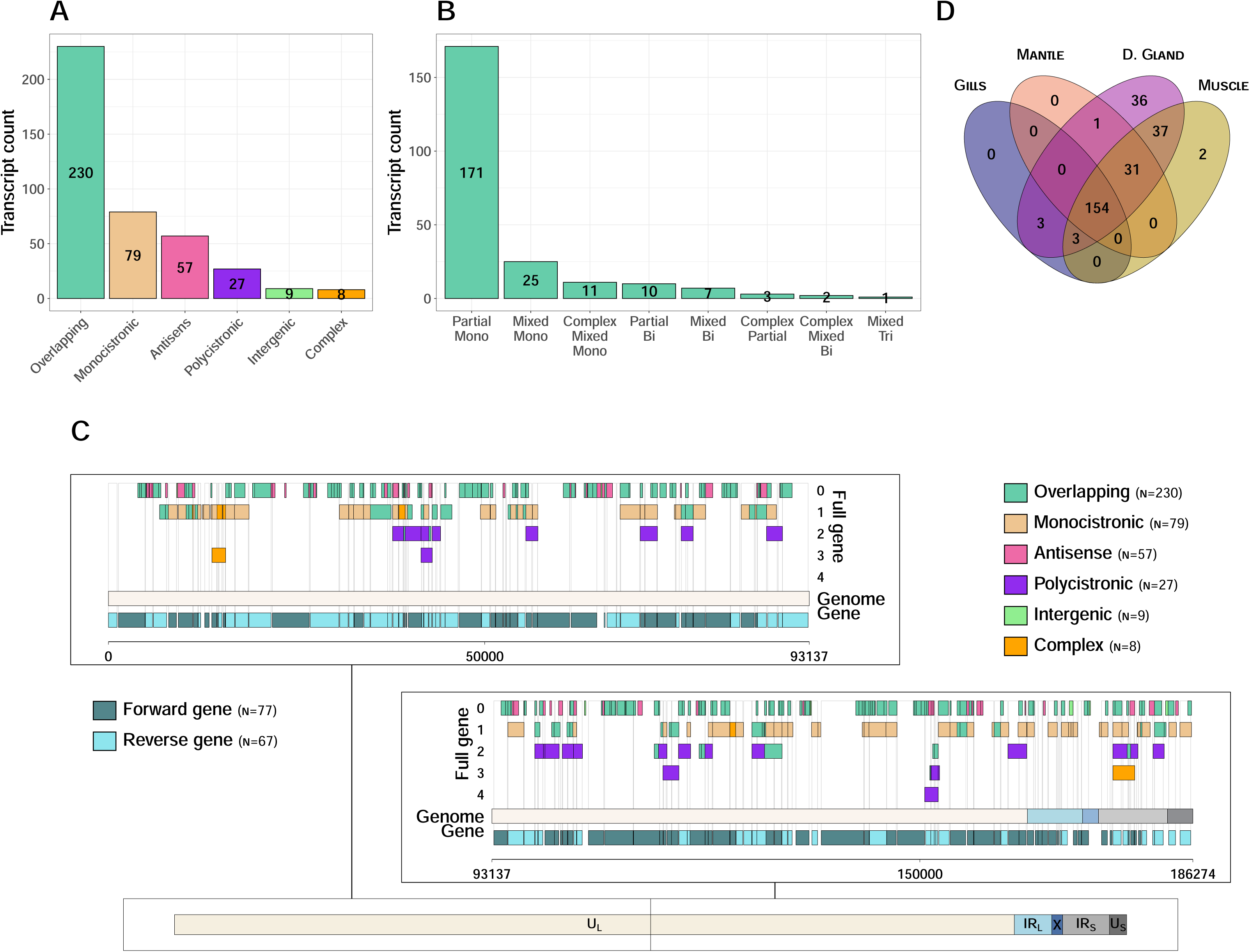
Novel viral transcript characterization. A. Histogram showing the six transcript categories. The y-axis shows transcript counts and the x-axis shows the six categories. B. Histogram showing the six subtypes within the ’overlapping’ category. The y-axis shows transcript counts and the x-axis shows the subtypes. C. LoRTIA-identified transcripts aligned to the OsHV-1 genome. The panel is divided into two subpanels: the top subpanel displays the first half of the genome, whereas the bottom subpanel displays the second half. Transcripts are plotted along the genome and colored according to the category to which they belong. They are positioned on the y-axis according to the number of full genes they overlap. Annotated OsHV-1 genes on the forward and reverse strands are shown at the bottom of each subpanel. The x-axis represents genomic position in base pairs. D. This venn diagram illustrates the number of transcripts with expression levels greater than 0 TPM across various tissues, considering all infection levels

The transcripts were then compared with the 144 annotated OsHV-1 ORFs (Figure 6.C, Table S4). No spliced transcript was detected. Of the 144 ORFs, 102 were entirely covered by at least one sense transcript and 27 were only partially covered, including those encoding the putative DNA primases (ORF49 and ORF66), DNA helicase (ORF67) and DNA polymerase (ORF100). Five ORFs were overlapped only by antisense transcripts (ORF5.1, ORF50.1, ORF109, ORFS43.3 and ORFS72.1). Ten ORFs, none with a predicted function, were not overlapped by any transcript (ORF1, ORF6, ORF91, ORF5.2, ORF50.2, ORFS4.1, ORFS4.2, ORFS115.2, ORFS116.1 and ORFSIN1.2). Moreover, three transcripts were also detected within the X region. Two of them partially overlapped ORFS115 and one was intergenic. None of them extended across the IR_L_-X or X-IR_S_ boundaries.

#### 4.2. OsHV-1 transcript expression varied mainly with infection level

A redundancy analysis was performed on Hellinger-transformed proportions of the 410 OsHV-1 transcripts across the 36 libraries from infected oysters, with infection level and tissue as explanatory variables (Figure S4.A). Together, these two factors explained 26.9% of the total variance (adjusted R² = 14.7%). The first constrained axis (RDA1) accounted for 68.02% of the constrained variance and separated libraries according to infection level, which had a significant effect on transcript composition when permutations were restricted to whole oysters (F = 4.25, df = 2, p = 0.0047). The second axis (RDA2) accounted for 12.83% of the constrained variance; tissue did not have a significant effect once permutations were restricted within oyster (F = 0.84, df = 3, p = 0.921), despite the apparent clustering by tissue visible on the ordination plot.

#### 4.3. OsHV-1 transcript detection and abundance are shaped by transcript length and category, independently of tissue or infection level

Of the 410 OsHV-1 transcripts, 154 (37.6%) were detected in all four tissues (detection criterion: ≥5 normalized counts in at least 2 libraries per tissue) (Figure 6.D and Table 1); the remaining 256 showed partial (n = 113) or no detection (n = 143). Partially or non-detected transcripts were markedly less expressed than those detected in all four tissues (median maximum expression: 15.3 vs 299 normalized counts; Table S4). This pattern was not attributable to genuine tissue specificity. Indeed, a permutation test at equalized sequencing depth showed no difference from random tissue assignment (409/410 observed vs. a median of 409/410 expected, p = 1), consistent with a detection-limit explanation.

**Table 1:** Detection completeness and transcript length by OsHV-1 transcript category.

| **Category** | **Total transcripts** | **Detected in all 4 tissues** | **Detection rate (%)** | **Median length (bp)** |
|---|---|---|---|---|
| Intergenic | 9 | 5 | 55.6 | 191.0 |
| Monocistronic | 79 | 37 | 46.8 | 944.0 |
| Overlapping | 230 | 99 | 43.0 | 703.0 |
| Antisense | 57 | 11 | 19.3 | 373.0 |
| Complex | 8 | 1 | 12.5 | 1 451.5 |
| Polycistronic | 27 | 1 | 3.7 | 1 495.0 |
| **Total** | **410** | **154** | **37.6** | **-** |

Detection completeness also varied by transcript category (χ² = 30.5, df = 5, p < 0.0001), with polycistronic (3.7%), complex (12.5%) and antisense (19.3%) transcripts showing the lowest detection rates across all four tissues, compared to intergenic (55.6%), monocistronic (46.8%) and overlapping (43.0%) transcripts (dataset-wide average: 37.6%; Table 1). This pattern was largely explained by transcript length. Transcripts detected in all four tissues were shorter than partially or non-detected ones (median 536 vs 936 bp, p = 8 × 10⁻¹⁵), and length correlated negatively with the number of tissues detected (Spearman’s ρ = -0.40, p < 2.2 × 10⁻¹⁶). Logistic regression confirmed that length and category contributed independently to detection completeness (both p < 10⁻⁹ when added to the other). After accounting for length, antisense transcripts remained associated with a higher probability of non-detection than other categories (p < 0.001 for monocistronic and overlapping vs. antisense), whereas the apparent under-detection of polycistronic transcripts was substantially attenuated once length was taken into account (Table 1).

Given the low detection completeness of polycistronic transcripts, their abundance was further examined directly against monocistronic transcripts sharing the same ORF. Sixteen ORFs were represented in both polycistronic and monocistronic transcripts (16 monocistronic and 15 polycistronic transcripts). Across these ORFs, polycistronic transcripts were less abundant than their monocistronic counterpart (median log2 ratio = -1.59, i.e. ∼3-fold lower; Wilcoxon signed-rank test, p = 1.5 × 10⁻⁴), with lower polycistronic expression for 14 of the 16 ORFs (Figure S5). Polycistronic transcripts were more abundant in highly than in moderately infected oysters, for all 16 ORFs (median fold-change = 1.9×), but this increase was smaller than for monocistronic transcripts (median fold-change = 3.6×; paired Wilcoxon test, p = 0.0042).

#### 4.4. Few OsHV-1 transcripts showed tissue-dependent abundance

Differential expression of OsHV-1 transcripts between tissues was tested separately for each infection level and for each pair of tissues (18 comparisons, DESeq2, adjusted p < 0.05, |log2FC| > 2) (Table S4). No transcript was differentially abundant between tissues in lightly or moderately infected oysters. In highly infected oysters, three transcripts were more abundant in the gills or mantle relative to the digestive gland. An overlapping transcript associated with ORF10 (no predicted function or domain; tr17) was more abundant in both the gills (log2FC = 2.19, padj = 0.044) and the mantle (log2FC = 4.04, padj = 0.011) than in the digestive gland. A polycistronic transcript associated with ORF73.1 and ORF73.2 (no predicted function or domain; tr345, log2FC = 2.73, padj = 0.039) and a monocistronic transcript associated with the RING-type zinc-finger gene ORF118 (tr602, log2FC = 2.42, padj = 0.044) were also more abundant in the mantle than in the digestive gland (Table 2). These three transcripts spanned different categories and genomic regions, with no shared functional annotation. No other tissue pair showed differentially abundant transcripts at any infection level.

**Table 2:** OsHV-1 transcripts differentially abundant between tissues in highly infected oysters.

| **Transcript** | **Comparison** | **Log2FC** | **padj** | **Category** |
|---|---|---|---|---|
| tr17 | Gills vs D. Gland | 2.19 | 0.044 | Overlapping |
| tr17 | Mantle vs D. Gland | 4.04 | 0.011 | Overlapping |
| tr345 | Mantle vs D. Gland | 2.73 | 0.039 | Polycistronic |
| tr602 | Mantle vs D. Gland | 2.42 | 0.044.3 | Monocistronic |

### 5. Host transcriptional response to OsHV-1 infection

#### 5.1. Host transcript expression varied mainly with tissue and secondarily with infection level

RDA analysis was applied to the vst counts of the *M. gigas* transcripts in the 48 libraries, with tissue and infection level as explanatory variables (Figure S3.B). Together, these two factors explained 87.3% of the total variance (adjusted R² = 0.854). The first constrained axis (RDA1) accounted for 52.08% of the constrained variance and separated libraries according to tissue. The second axis (RDA2) accounted for 23.84% and separated libraries according to infection level. Both tissue (permutations restricted within oysters; F = 79.7, p < 10⁻⁴) and infection level (permutation of whole oysters; F = 13.9, p < 10⁻⁴) had significant effects.

#### 5.2. Two host co-expression modules were associated with OsHV-1 infection

Weighted gene co-expression network analysis (WGCNA) identified ten co-expression modules (Figure S6.A and Table S5). Two of them met the selection criterion (p < 0.01, r² > 0.5) for a positive correlation with both infection level and viral load, namely the brown module (3,343 transcripts; r = 0.83 and 0.85, respectively) and the magenta module (202 transcripts; r = 0.56 and 0.59, respectively). These correlations remained significant in linear mixed models on module eigengenes with oyster as a random effect (brown: p = 3.0 × 10⁻¹² for infection level, p = 3.1 × 10⁻⁵ for viral load; magenta: p = 2.4 × 10⁻⁴ for infection level, p = 7.6 × 10⁻⁵ for viral load), meaning that the observed associations are not driven by non-independence among the four tissues sampled from the same oyster.

The brown module was enriched in 135 biological process, 43 cellular component and 22 molecular function GO terms (Table S6). These terms were mainly related to RNA processing and splicing, regulation of transcription, DNA damage response, cellular stress response, protein phosphorylation, proteolysis and apoptosis, and vesicle-mediated transport and Golgi organization. The magenta module was enriched in 39 biological process, 9 cellular component and 15 molecular function terms. These were mainly related to cellular stress response, endoplasmic reticulum stress and regulation of cell survival.

The brown module eigengene increased with infection level in all four tissues (Figure S6.B), with no significant tissue × infection level interaction (linear mixed model with oyster as random effect; F = 0.49, df = 3, p = 0.69). The magenta module eigengene, in contrast, increased sharply with infection level in the mantle and more gradually in the gills, and varied little in the digestive gland and muscle, reflected in a significant tissue × infection level interaction (F = 11.8, df = 3, p = 2.8 × 10⁻⁵) (Figure S6.B).

#### 5.3. The host response to infection was strongest in the gills and mantle

To compare the host response among tissues, differential expression analysis was performed separately for each tissue. Lightly, moderately and highly infected oysters were each compared with not infected oysters (Figure 7.A, Table S5). The number of upregulated differentially expressed transcripts (DETs) increased with infection level in the mantle, digestive gland and muscle (lightly infected: 3-86; moderately infected: 16-87), but not in the gills, where upregulated DETs were already numerous at the lightly infected stage (86) and remained at a similar level in moderately infected oysters (87). In highly infected oysters, upregulated DETs were more numerous in the gills (268) and mantle (236) than in the muscle (95) and digestive gland (40).

**Figure 7:**
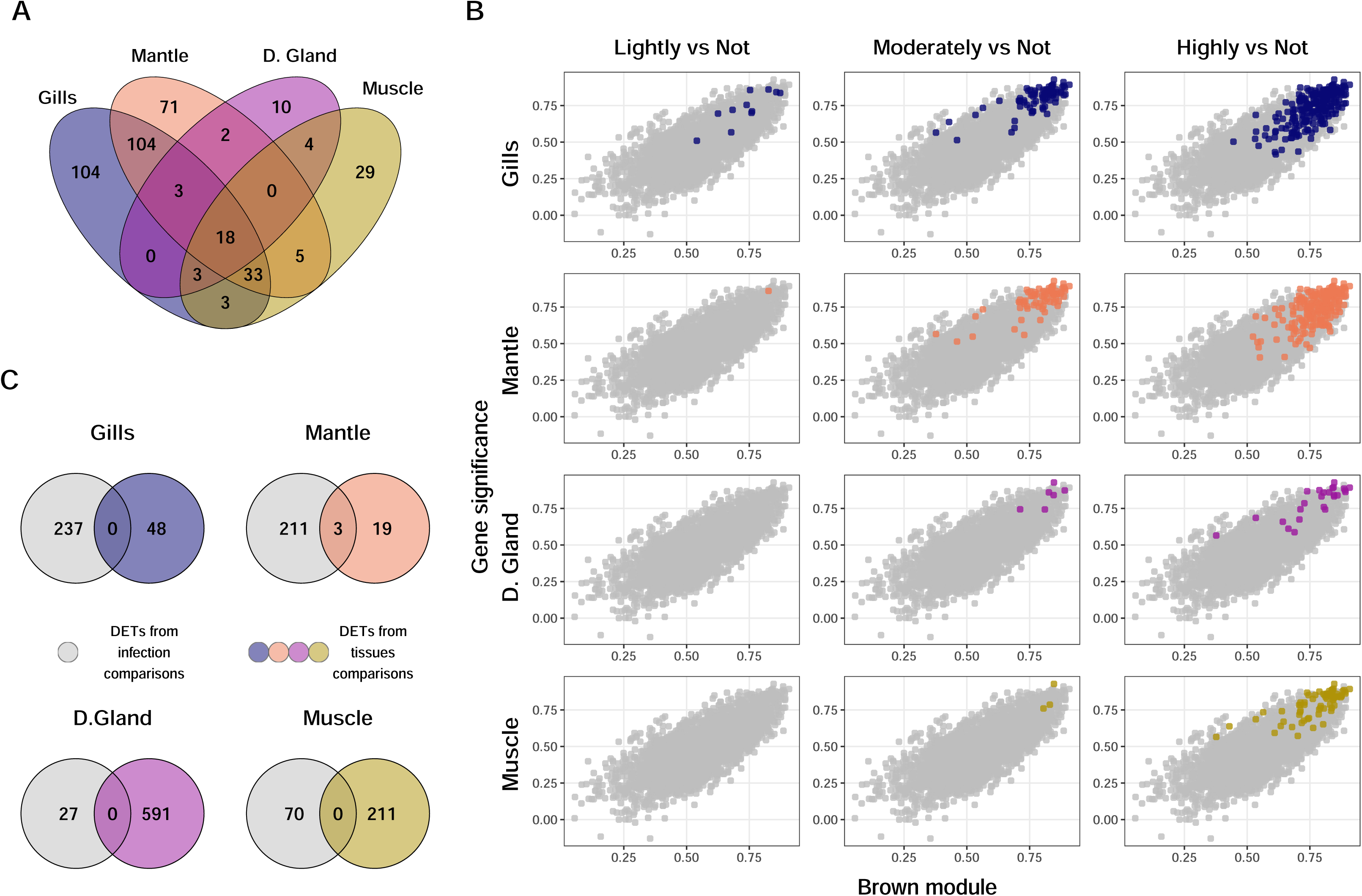
Oyster DETs analysis. A. Venn diagram of oyster DETs in the highly infected condition B. DETs within the brown module. The Y-axis shows the gene significance values obtained from the WGCNA analysis, and the X-axis shows the module membership scores for the brown module. Each dot represents a transcript assigned to this module. The plot highlights the DETs that were found to be upregulated by DESeq2 across different tissues (rows) and infection levels (columns). Colors indicate tissue types. C. Venn diagram of DETs shared with the brown module and DETs identified from tissues comparison (at highly infected condition).

Most upregulated DETs of highly infected oysters belonged to the brown module: 81.3% in the gills, 83.1% in the mantle, 70.5% in the muscle and 62.5% in the digestive gland. Between 3.2% and 6.4% belonged to the magenta module. From lightly to highly infected oysters, the number of brown module DETs increased from 8 to 218 in the gills, from 1 to 196 in the mantle, from 0 to 67 in the muscle and from 0 to 25 in the digestive gland, with intermediate counts of 75, 72, 3 and 6 respectively in moderately infected oysters (Figure 7.B). Within each tissue, the brown module eigengene was correlated with the proportion of OsHV-1 reads in the library (Spearman’s r = 0.63–0.88, highest in the mantle).

#### 5.4. Part of the host response to infection was specific to the mantle

To identify tissue-specific features of the host response, the four tissues of highly infected oysters were compared pairwise using a paired design accounting for individual oysters. Transcripts differentially expressed in a single tissue relative to each of the three other tissues were retained. In total, 48 transcripts were specific to the gills, 22 to the mantle, 591 to the digestive gland and 211 to the muscle. Most of them belonged to modules correlated with tissue but not with infection (Table S5), and therefore mainly reflected constitutive differences between tissues.

Three of the 22 mantle-specific transcripts belonged to the magenta module and were also differentially expressed between highly and not infected oysters within the mantle. These transcripts showed a mantle-specific pattern, with markedly higher abundance in the mantle than in the digestive gland (log2FC = 5.3 to 6.6) and muscle (log2FC = 6.1 to 8.7). This overlap was not attributable to greater host sequencing depth in the mantle, whose median host read count was intermediate among the four tissues (Table S5). They corresponded to three genes: two kappaPI-actitoxin-like genes (LOC105339868, LOC136270124) and surfeit locus protein 2 (LOC105327470). No gill-specific or muscle-specific transcript met this combined criterion, nor did any digestive-gland-specific transcript (Figure 7.C).

## DISCUSSION

Despite increasing knowledge on OsHV-1 life cycle, the mechanisms regulating its lytic phase, the functional relevance of its genomic isomers, and the complexity of its transcriptome during *in vivo* infection remain poorly understood. Our study aimed to improve our understanding of OsHV-1 lytic phase by applying long-read sequencing across multiple Pacific oyster tissues in the context of an experimental infection. To mimic natural infection conditions, oysters were infected *via* bath exposure using contaminated seawater. While this design simplifies the environmental complexity of field settings, it is well suited to decipher OsHV-1 life-cycle dynamics and host responses (Agnew et al., 2020; Burge et al., 2020; de Lorgeril et al., 2018).

In the infected control tank, 64% of oysters died during the experiment, with mortality beginning 78 hours post-infection (hpi). Moribund oysters from all tanks exhibited high levels of viral DNA, confirming that mortality was associated with OsHV-1 infection. At the onset of the experiment, seawater contained approximately 2 × 10³ OsHV-1 genome copies per µL. Given a tank volume of approximately 35 L, this corresponds to a total of approximately 7 × 10¹⁰ genome copies in the tank. As the qPCR assay used cannot discriminate between infectious and non-infectious viral particles, nor between encapsidated viral genomes and free viral DNA, this value represents an upper bound on the number of infectious viral particles present. Assuming a homogeneous distribution of viral particles in the tank and an average filtration rate of 0.5 L of seawater per hour for juvenile oysters (Bougrier et al., 1995; Schikorski, Faury, et al., 2011), each oyster would have been exposed to approximately 10⁹ viral genome copies per hour at the onset of the experiment. This theoretical exposure is well above the LD50 estimated by intramuscular injection, which is only a few hundred undamaged viral capsids (Renault et al., 2024). Although exposure through filtration and injection are not directly comparable, as not all filtered particles are retained and the viral concentration in seawater changes over time, this suggests that oysters were exposed to a viral dose largely sufficient to initiate infection, consistent with the mortality observed.

The dynamics of OsHV-1 genome copy number in oysters and seawater indicate that viral particles were taken up from seawater within 3 hpi, as indicated by the sharp decrease in seawater viral DNA, followed by an active replication phase lasting approximately 50 hours. This phase preceded the release of viral particles into the surrounding seawater, with viral DNA in seawater reaching a maximum around 72 hpi, just before the onset of oyster mortality.

Furthermore, substantial variability in viral load was observed among oysters, indicating that individuals were at different stages of infection, as previously reported for OsHV-1 (Burge et al., 2020; Sauvage et al., 2009). Since all oysters originated from the same cohort and were exposed to the same seawater, this variability is likely to reflect host-related factors, among which filtration activity may play a role. OsHV-1 is transmitted horizontally through seawater (Schikorski, Faury, et al., 2011), and viral detection in epithelial cells of organs in contact with the external environment, such as the gills and mantle, suggests that these tissues may act as portals of entry for the virus (Lipart & Renault, 2002; Martenot et al., 2016). The amount of virus encountered by each oyster is therefore likely to depend on the volume of seawater it filters. Filtration rate varies substantially among individual oysters depending on size, valve activity and physiological state (Ehrich & Harris, 2015), and oyster size has itself been shown to affect susceptibility to OsHV-1 (Dégremont, 2013). Oysters exposed to the same viral concentration are thus unlikely to have filtered equivalent amounts of virus.

Due to cost constraints, only 12 oysters were selected for long-read genomic and transcriptomic sequencing to investigate the lytic phase of OsHV-1. Although this represents a small fraction of the total experimental population, the selected individuals were chosen to reflect the observed replication dynamics. In the lightly infected condition, only a small number of OsHV-1 genomic sequences were detected, covering only a limited portion of the viral genome and accompanied by low levels of viral gene expression. This stage corresponds to a viral load of approximately 10 genome copies per ng of DNA, consistent with the average load measured by qPCR in oyster tissues collected before 25 hpi. These findings suggest that OsHV-1 could be in the pre-replicative phase prior to genome replication. By contrast, in moderately infected conditions, the whole OsHV-1 genome was covered by genomic sequences and gene expression levels were significantly higher. This stage corresponds to a viral load of around 10³ genome copy numbers per ng of DNA, which matches the average viral load measured between 50 hpi and 75 hpi. These observations suggest that viral replication was fully established at this stage. Finally, in highly infected conditions, both OsHV-1 genomic and transcriptomic sequences were markedly elevated. This condition corresponds to a viral load of around 10⁵ genome copy numbers per ng of DNA, which reflects the high level of infection usually seen at the end of the OsHV-1 infection cycle (Sauvage et al., 2009; Schikorski, Faury, et al., 2011).

The fraction of reads aligning to neither reference genome was also characterized to verify that no other agent could confound the interpretation of these data (Supplementary File 1). It was dominated by bacterial sequences and by a single ciliate species, with no other viral agent detected that could plausibly have contributed to the mortality observed. The only RNA viruses recovered were several distinct picorna-like viruses (order *Picornavirales*) related to viruses recently identified in Pacific oysters from British Columbia (Zhong et al., 2026), detected almost exclusively in the digestive gland and present at all infection levels, including in uninfected oysters. This tissue-restricted, infection-independent pattern is consistent with the accumulation of environmental or dietary viruses through filter feeding rather than with an infection of the oyster itself, as repeatedly proposed for RNA viruses recovered from bivalve transcriptomes (Rosani et al., 2019; Wu et al., 2024). None of these sequences is therefore likely to interfere with the interpretation of the OsHV-1 data.

One of the aims of this study was to characterize the genomic plasticity of OsHV-1 during its lytic phase across different tissues of the Pacific oyster. To this end, the first step was to assemble a complete new OsHV-1 genome specific to this study. This *de novo* assembled genome was found to be most similar to a previously characterized viral genome isolated from Pacific oysters originating from the same geographical area and collected within a similar timeframe (Dotto-Maurel, Pelletier, et al., 2025), consistent with the reported spatiotemporal structuring of OsHV-1 genomic diversity (Delmotte et al., 2022; Morga et al., 2017).

To assess whether viral genetic diversity exhibited tissue-specific structure, we compared the distribution of intra-host SNPs across tissues. SNP profiles were consistently shared across tissues within each individual oyster, with no evidence of tissue-restricted variants, indicating that the OsHV-1 population remains genetically homogeneous across infected tissues during the lytic phase. This finding is consistent with studies on human herpesviruses, in which viral genomes recovered from multiple organs of the same individual were largely conserved, suggesting limited genetic structuration among tissues (Pyöriä et al., 2024; Santiago et al., 2021). Those studies, however, examined persistent infections lasting several years, whereas our results show that this homogeneity is already established within a single acute lytic episode. By contrast, the number and identity of detected variants differed substantially among oysters, suggesting that the host, rather than tissue type, is the main source of intra-host viral genetic diversity. This is in line with a recent preprint on OsHV-1, which similarly proposed that host-to-host variation and associated selective pressures are the primary driver of viral diversity (Pelletier et al., 2026).

Then, upon realigning the OsHV-1 genomic sequences to the newly assembled genome, we detected the four major genomic isomers in all tissues except muscle. This observation may be explained by the fact that OsHV-1 genomic sequences obtained from muscle tissue were shorter than those from other tissues, suggesting that the absence of detectable isomer reads in muscle could be due to the reduced length of sequenced DNA fragments in this tissue. As previously reported, we also identified reads corresponding to the minor isomers T_X_ and N_X_ (which lack the X region between IR_L_ and IR_S_) in the gills and mantle, but not in the digestive gland (Dotto-Maurel et al., 2022; Dotto-Maurel, Pelletier, et al., 2025). However, as for muscle, this observation may be attributed to the shorter length of sequenced DNA fragments in this tissue. While the relative proportions of the four major isomers appeared similar across different tissues, the overall number of long reads in each condition was insufficient for the complete characterization of these isomers or definitive conclusions about their tissue specificity. Improving the recovery of high-molecular-weight DNA, ideally yielding full-length OsHV-1 genomes as single DNA molecules, would substantially improve the characterization of genome isomerization. Interestingly, although long DNA reads provided evidence for the presence of the minor N_X_ and T_X_ isomers, no cDNA reads mapped to the IR_L_-IR_S_ junction, and no OsHV-1 transcripts were detected in this region. This absence of detectable transcription suggests that these minor isomers are unlikely to play a direct role in viral gene expression during infection in oysters. Instead, their occurrence is consistent with the hypothesis proposed for other herpesviruses that genome isomers arise as by-products of the viral DNA replication process rather than representing biologically distinct transcriptional states (Jacob et al., 1979; Weber et al., 1988).

Our next aim was to characterize the dynamics of OsHV-1 gene expression during its lytic phase in different oyster tissues. While there were differences in expression levels, the overall viral gene expression profiles were highly similar among tissues, with the highest coverage consistently observed in the mantle. The five most highly expressed viral genes were consistently ORF18 (predicted intrinsically disordered protein), ORF27 (dUTPase-like protein), ORF80 (predicted transmembrane protein), ORF107 (predicted intrinsically disordered protein), and ORF120 (predicted membrane protein). This expression pattern closely mirrors results previously reported in OsHV-1-infected haemocytes maintained *in vitro*, where ORF80, ORF27 and ORF107 also ranked among the top five most highly expressed genes (Dotto-Maurel, Le Luyer, et al., 2025). Moreover, ORF27 was identified as highly expressed in the mantle of Pacific oysters (Segarra et al., 2014), proteins coded by ORF107 and ORF27 have also been identified as highly present in Pacific oysters infected by OsHV-1 (Leprêtre et al., 2021), and ORF107 was among the most abundant genes expressed in the OsHV-1 isolate ZK0118 infecting clam hemolymph (Rosani et al., 2024). Furthermore, viral genes were grouped into three distinct expression clusters, consistent with the clustering previously reported in the *in vitro* haemocyte study (Dotto-Maurel, Le Luyer, et al., 2025). As observed in that study, the five most highly expressed genes belonged to the transcriptional cluster associated with the lightly infected condition, suggesting that they are induced at the onset of infection. The conservation of transcriptional clustering and expression kinetics in our *in vivo* dataset compared to previously published *in vitro* data suggests that the temporal organization of the OsHV-1 lytic transcriptional program is preserved across these two infection models. The abundant expression of these five genes in the lightly infected condition further suggests that they play important roles during lytic infection, despite their functions remaining largely unknown. Nevertheless, the predicted structural features of their coded proteins offer valuable insights into their potential functions. ORF27 codes for a dUTPase-like enzyme, which in other herpesviruses plays a critical role in maintaining genome integrity by preventing misincorporation of uracil during viral DNA replication (Kato et al., 2014). The early and high-level expression of ORF27 is therefore consistent with a role in supporting viral DNA replication from the very onset of infection. ORF80 codes for a transmembrane glycoprotein predicted to localize to the Golgi apparatus (Dotto-Maurel, Le Luyer, et al., 2025), a compartment central to virion envelopment in herpesviruses. Although herpesvirus glycoproteins are primarily associated with virion assembly, entry, and cell-to-cell spread (Harkness et al., 2014; Knipe et al., 2013; Volkening et al., 2023), envelope glycoproteins can also exert regulatory functions by modulating membrane fusion, intracellular signaling, or host-virus interactions (Connolly et al., 2011). Therefore, the early expression profile of ORF80 raises the possibility that this glycoprotein may contribute to additional regulatory processes during the early stages of the lytic cycle. ORF18, ORF107, and ORF120 remain entirely uncharacterized to date. Notably, both ORF18 and ORF107 code for proteins containing predicted intrinsically disordered regions, which are known to mediate flexible interactions with multiple binding partners and are frequently involved in transcriptional regulation, signaling pathways, and virus-host interactions (Trivedi & Nagarajaram, 2022). Their early and high expression at the lightly infected stage therefore suggests that they may play important regulatory roles during the initiation of the OsHV-1 lytic cycle.

One of the key advantages of long-read sequencing is its ability to capture full-length transcripts. Here, we present the first long-read transcriptome of OsHV-1 generated from infected Pacific oyster tissues. Previous long-read studies of OsHV-1 were restricted to the genotype infecting clams, which does not infect *M. gigas* (Bai et al., 2021; Rosani et al., 2024), whereas transcriptomic data available for *M. gigas* relied on short reads, which fragment transcription units. The 410 transcripts identified here provide a transcript-level annotation of the viral genome in its natural host. No spliced transcript was detected, and only 79 transcripts (19%) fully covered a single ORF and were classified as monocistronic. Although part of the remaining transcripts likely reflects incomplete reads or degradation, the presence of fully covered antisense (n = 57) and polycistronic (n = 27) transcripts shows that OsHV-1 shares with other herpesviruses a transcriptional architecture more complex than its ORF annotation suggests (Boldogkői, Moldován, et al., 2019; Tombácz et al., 2019). Of the 144 annotated ORFs, 134 were overlapped by at least one transcript, whereas no transcript was detected for 10 of them, none of which have a predicted function. Several of these genes were previously reported as not being expressed in Pacific oyster infected haemocytes maintained *in vitro* (ORFS4.2 and ORFS116.1) and in blood clam haemolymph infected *in vivo* (ORF1 and ORF5.2) (Dotto-Maurel, Le Luyer, et al., 2025; Rosani et al., 2024). Their absence from the reconstructed transcriptome therefore likely reflects a lack of expression under the experimental conditions used here, expression restricted to another stage of the viral life cycle, or inaccuracies in the current genome annotation. Additionally, transcripts could only be partially reconstructed for 27 annotated genes. To identify transcript boundaries, we used LoRTIA, which detects transcription start and end sites from long-read sequencing data using adapter- and end-based evidence. This approach has been extensively validated in herpesvirus transcriptomes (Boldogkői et al., 2018; Maróti et al., 2021; Prazsák et al., 2024; Tombácz et al., 2025; Torma et al., 2023, 2025) and is well suited to compact viral genomes, where transcript diversity mainly results from alternative transcription initiation and termination rather than extensive splicing. Manual inspection of read alignments confirmed that LoRTIA-defined boundaries were consistent with coverage patterns.

For the 27 partially supported genes, incomplete transcript reconstruction was mainly due to the absence of either the 5′ or 3′ adapter required for full-length detection. Combined with low read coverage in these regions, this likely reflects technical limitations of the adapter-dependent approach rather than a lack of transcription. Notably, four genes involved in DNA replication (ORF49, ORF66, ORF67 and ORF100) were among the 27 partially supported genes. These genes were expressed at low levels both in our experiment and in previous studies of infected haemocytes maintained *in vitro* and in haemolymph collected from infected clams (Bai et al., 2021; Dotto-Maurel, Le Luyer, et al., 2025). Their consistently low expression suggests that these transcripts may be required only at relatively low abundance during OsHV-1 genome replication. Increasing sequencing depth and library construction efficiency could enable the recovery of reads that support the expression of these genes, which contain adapters at both ends. This would clarify the organization of transcripts in these regions.

Detection completeness of OsHV-1 transcripts across tissues was strongly associated with transcript length, independently of transcript category. This pattern is consistent with the known bias of PCR-based long-read cDNA sequencing towards shorter molecules, which can reduce the representation of longer transcripts in the final library independently of their true abundance (Bayega et al., 2022; Shagin et al., 1999). This bias is particularly relevant for polycistronic transcripts, which were the longest and least completely detected category and may therefore partly account for their reduced apparent abundance relative to monocistronic transcripts sharing the same ORF. However, the persistent difference between the two transcript categories, which became more pronounced with increasing infection level, suggests that technical bias alone is insufficient to explain the observed pattern. This result is consistent with observations in other herpesviruses, where polycistronic transcripts are commonly found but typically expressed at lower levels than their monocistronic counterparts (Boldogkoi, 2012; Boldogkői, Tombácz, et al., 2019; Tombácz et al., 2017, 2019). While their precise function remains unclear, it has been suggested that such transcripts may play a role in gene regulation by interfering with, or modulating, the transcription of neighboring genes (Boldogkői, Tombácz, et al., 2019; Tombácz et al., 2019). Moreover, antisense transcripts also remained under-detected after accounting for length, indicating either an additional category-specific bias or genuinely lower expression. The large number of overlapping transcripts identified here may further include degradation products that meet the criteria for full-length transcripts, although distinguishing these from genuine short isoforms will require complementary approaches such as direct RNA sequencing or proteomic validation (Prawer et al., 2023).

In contrast, tissue had little influence on transcript detection or abundance. A permutation test showed that incomplete detection across tissues did not differ from random expectations after equalizing sequencing depth, while compositional analysis revealed no significant tissue effect once individual oysters were accounted for. Only three transcripts differed in abundance between tissues in highly infected oysters, with no clear functional or genomic pattern. One of the transcripts more abundant in mantle than digestive gland, tr602, encodes a RING-type zinc-finger protein (ORF118), a domain associated with E3 ubiquitin ligase activity and immune modulation in other herpesviruses (Boutell et al., 2002; Boutell & Everett, 2013; R. D. Everett et al., 2010; Hagglund et al., 2002). Overall, these results suggest that OsHV-1 transcription is largely conserved across host tissues, with transcript length and category exerting substantially greater influence on detection and abundance than tissue.

Host transcriptomic responses to OsHV-1 infection, by contrast, varied more clearly across tissues. The gills and mantle displayed the largest number of upregulated transcripts in response to infection, most of which belonged to the brown co-expression module, whose eigengene increased with infection level uniformly across all four tissues. The digestive gland and muscle showed markedly fewer differentially expressed host transcripts, suggesting a comparatively weaker transcriptional response in these tissues.

The functional enrichment analysis of the infection-associated brown module provides insight into the molecular functions, biological processes, and cellular components represented within this co-expression network, thereby helping to identify host pathways and cellular systems associated with infection. Several enriched GO terms point to processes that are commonly associated with host-virus interactions. Terms related to apoptosis and DNA damage response are consistent with classical antiviral strategies aimed at eliminating infected cells, although they could equally reflect virus-driven manipulation of host cell death pathways to favor its own replication (H. Everett & McFadden, 1999; Yatim & Albert, 2011). Similarly, the enrichment in RNA processing, splicing, and transcriptional regulation may indicate either a host-driven reprogramming of gene expression in response to infection or, conversely, a viral strategy to hijack host transcription machinery (Liao & Garcia-Blanco, 2021). Of particular interest is the enrichment in vesicle-mediated transport and Golgi organization, given that herpesviruses are known to rely on the host secretory pathway for virion envelopment and egress (Dotto-Maurel, Arzul, et al., 2025; R. D. Everett, 2014; Knipe et al., 2013). This may reflect either host restriction mechanisms targeting this pathway or its exploitation by the virus.

This overall tissue gradient in response magnitude, however, does not extend to which tissue-specific transcripts fall within infection-associated modules and are themselves differentially expressed with infection. Tissue-specific transcripts in the digestive gland, gills and muscle showed no such overlap, consistent with constitutive, tissue-defining expression differences unrelated to infection. The mantle was the exception showing a marked mantle-specific abundance pattern. Two of the three corresponding genes were annotated by homology to kappaPI-actitoxin-like sequences, a family of secreted peptides characterized in sea anemones as serine protease inhibitors with reported toxin activity (da Silva et al., 2023; Mourão & Schwartz, 2013); whether this annotation reflects a genuinely analogous function in oyster immune defense, such as regulation of host or pathogen proteolytic cascades, remains speculative and would require functional validation.

Taken together, these results suggest that the OsHV-1 transcriptional program itself does not markedly differ between tissues, but that the host mounts a broad, magnitude-graded response in the gills and mantle, together with a tissue-specific, infection-linked signature concentrated in the mantle. This is consistent with previous reports pointing to pallial organs, particularly the gills, as a primary site of OsHV-1 entry and activity (Arzul et al., 2002; Martenot et al., 2016; Sauvage et al., 2009; Segarra et al., 2016). The digestive gland and muscle, by comparison, showed comparatively limited host responses with little overlap with infection-associated modules, consistent with a role as less directly implicated tissues, or as recipients of secondary, non-specific physiological perturbation associated with systemic infection.

Beyond tissue-level patterns, these data refine our view of the OsHV-1 lytic cycle in four ways. First, the viral transcriptional program observed *in vivo* after natural-route exposure resembles that described in haemocytes *in vitro* (Dotto-Maurel, Le Luyer, et al., 2025). Indeed, three of the five most expressed genes are shared and viral genes fall into similar expression clusters. This suggests that the haemocyte model captures key features of the program operating in the whole animal, which is valuable in the absence of a permissive cell line. Second, this program appears largely tissue-invariant. Viral genome coverage, transcript composition and transcript abundance varied with infection level rather than with tissue, whereas the host response differed between tissues. The lytic cycle thus seems to proceed as a single, conserved program in every infected tissue, with tissue identity shaping mainly the host side of the interaction. Third, the gap between polycistronic and monocistronic transcripts widened as infection progressed, suggesting that transcript architecture changes along the lytic cycle. Although technical biases against long molecules may contribute, this pattern is compatible with a regulatory role of transcript overlaps, as proposed for other herpesviruses, and can now be tested experimentally. Finally, the structural plasticity of the viral genome, with the four major isomers detected in all tissues where long reads allowed it, had no detectable transcriptional counterpart, and intra-host genetic diversity was structured by individual oyster rather than by tissue. Because infection levels were used as a proxy for infection progression in a limited number of oysters, these conclusions concern the progression from early to advanced infection rather than fine-scale kinetics.

By combining a natural-route infection model with long-read sequencing of four tissues, this study provides the first long-read view of the OsHV-1 lytic cycle in *Magallana gigas* and a framework for refining viral genome annotation. Building on these data, further investigations combining microscopy, single-cell sequencing and proteomics will be needed to identify the cell types targeted by the virus, determine whether all infected tissues produce infectious particles, and elucidate the functional roles of the poorly characterized viral genes and of the polycistronic, complex and antisense transcripts identified here.

## MATERIALS & METHODS

The Materials and Methods are summarized in Figure 8.

**Figure 8:**
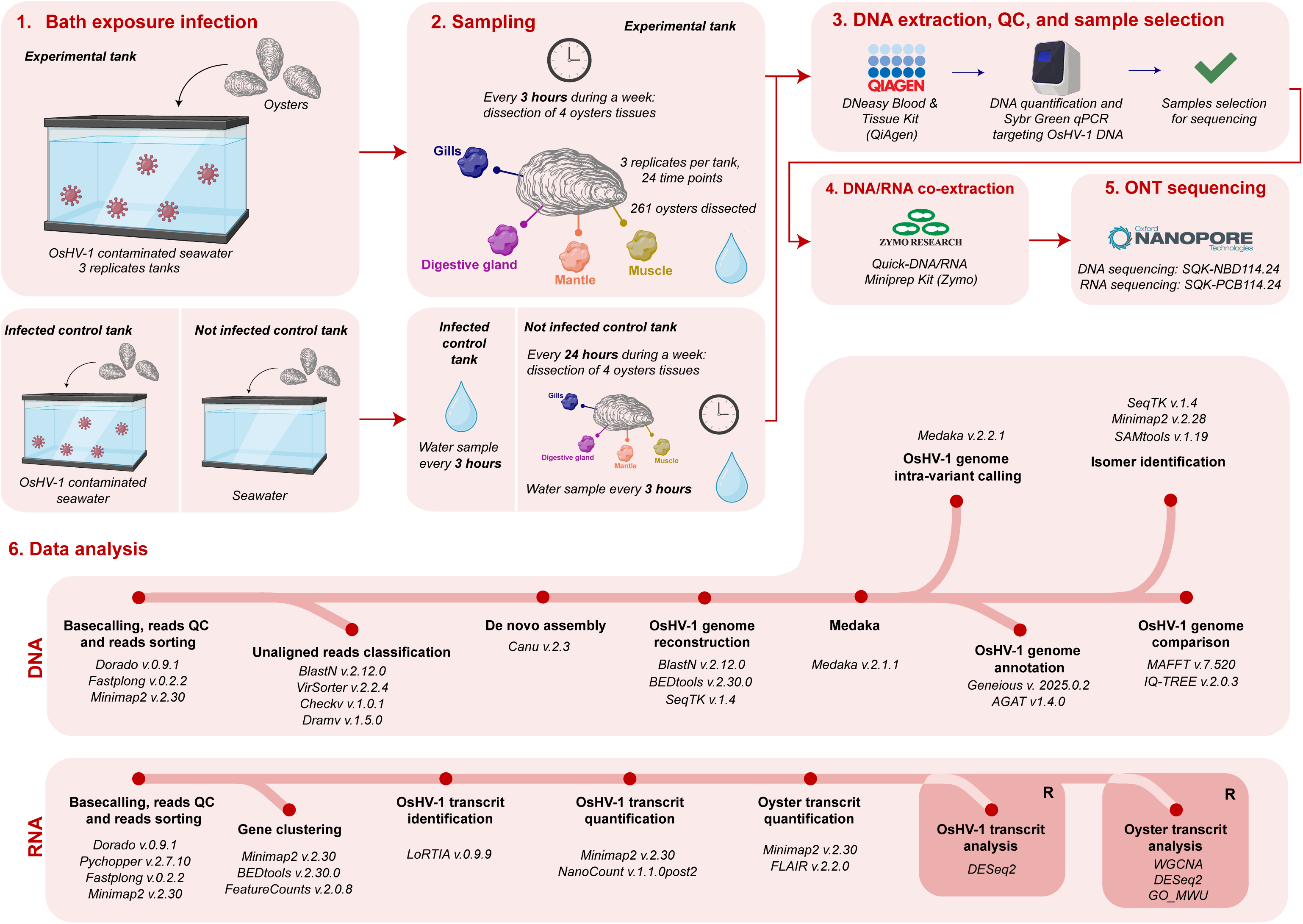
Experimental design of the study.

### 1. Experimental design

#### 1.1. Oysters production

Experimental oysters were produced from wild broodstock collected in Logonna-Daoulas (Rade de Brest, Brittany, France). Broodstock conditioning, gametogenesis induction, fertilization and larval rearing were performed following the protocol described previously (Dotto-Maurel et al., 2026). Briefly, broodstock was conditioned at 20°C in seawater enriched with a mixed microalgal diet (*Skeletonema costatum, Isochrysis galbana* and *Tetraselmis suecica*) to induce gametogenesis, and the resulting offspring were reared under controlled conditions in UV-treated seawater at the Ifremer facilities in La Tremblade. These oysters had therefore never been exposed to OsHV-1 prior to the experiment. Both donor and recipient oysters described below originated from this cohort. At the time of the experiment, they were nine months old, with a mean total weight of 20.5 g.

#### 1.2. OsHV-1 inoculum preparation

During a mortality event in August 2022, moribund oyster spat were collected from the La Floride oyster farming area (Marennes-Oléron Bay, France) (latitude 45.80° N, longitude -1.15°) and an initial viral inoculum was prepared as previously described (Schikorski, Renault, et al., 2011). In 2023, this inoculum was used to infect naive spat under laboratory conditions to produce sufficient volume for subsequent experiments, and a second inoculum was prepared from these animals following the same protocol. This second inoculum, corresponding to one laboratory passage of the original field isolate, was used in the present study.

#### 1.3. Production of OsHV-1 contaminated seawater

To characterize the lytic phase of OsHV-1 *in vivo,* OsHV-1 infection was performed by bath exposure. To this end we first generated contaminated seawater using donor oysters. To do so, 360 donor oysters were injected in the adductor muscle, using an AGANI™ needle (23G, 0.6×25mm), with 100 µL of a 1:10 diluted viral inoculum (1.28 × 10⁵ copies/µL), and distributed across three separate tanks, each containing 35 L of seawater (i.e., 120 oysters per tank). An infected control was also produced using 60 donor oysters infected with the same viral inoculum and placed in a separate tank containing 17 L of seawater. A not infected control was also included, consisting of 60 donor oysters injected with 100 µL of artificial seawater and housed in another tank with the same seawater volume.

Following injection, all tanks were maintained at 20°C for 42 hours to allow OsHV-1 replication and viral particles shedding into the tank seawater.

#### 1.4. Oyster infection and sampling

Forty-two hours after injection, the donor oysters were removed and replaced with 200 recipient oysters in each of the three experimental tanks and 100 recipient oysters in each of the two control tanks. This point marked the beginning of the infection experiment, defined as 0 hours post-infection (hpi).

Oysters were then sampled every 3 h over a 99-h period, with three individuals collected from each of the three challenged tanks at each time point. Gills, mantle, digestive gland and adductor muscle were dissected from each oyster, and two 20 mg subsamples were taken from each tissue. The first was stored dry at -20°C for DNA extraction (QIAamp DNA Kit, Qiagen) and qPCR quantification of OsHV-1 DNA; the second was stored at -20°C in 150 µL of DNA/RNA Shield (Zymo Research) for subsequent DNA/RNA co-extraction and Oxford Nanopore Technologies (ONT) sequencing.

Three oysters from the unexposed control tank were sampled and dissected every 24 h following the same procedure. In parallel, duplicate 1 mL seawater samples were collected from all tanks every 3 h and stored at -20°C prior to DNA extraction and qPCR quantification of viral DNA.

### 2. DNA extraction and OsHV-1 DNA quantification

One subsample of each dissected tissue and one of each seawater sample were used to quantify OsHV-1 genome copies over time. DNA was extracted in 96-well plates using the DNeasy Blood & Tissue Kit (Qiagen), and viral DNA was quantified by qPCR with the OsHVDP forward and reverse primers, as previously described (Webb et al., 2007). Each plate contained 90 samples analyzed as single technical replicates, a five-point standard curve generated from a plasmid carrying the target region (10¹ to 10⁵ copies per reaction) and a no-template control. Reactions were run on an Mx3005P thermal cycler and analysed with MxPro software (Agilent Technologies). Samples were considered positive for OsHV-1 DNA if a genuine amplification curve was obtained, along with a melting temperature consistent with that of the positive control.

### 3. Selection of samples for sequencing and tissues DNA/RNA co-extraction

On the basis of OsHV-1 DNA quantification, four dissected tissues were selected from three oysters per infection level, together with three oysters from the not infected control tank, for sequencing (n = 48 samples total; Table S1). Individual viral load was used as the selection criterion rather than sampling time, since infection progression was asynchronous both between individuals and between tanks, making viral load a more reliable indicator of infection stage than time elapsed since exposure. The corresponding sampling time point for each selected sample is reported in Table S1. Samples were classified as not infected, lightly infected (∼10 genome copies per ng DNA), moderately infected (∼10³ copies per ng DNA) or highly infected (∼10⁵ copies per ng DNA) (Table S1).

DNA and RNA were co-extracted from the second subsample of each selected sample using the Quick-DNA/RNA Miniprep Kit (Zymo Research) according to the manufacturer’s instructions, including the on-column DNase I treatment for the RNA fraction. Extracted RNA was split into three 20 µL aliquots to avoid repeated freeze-thaw cycles and stored at -80°C, while extracted DNA was stored at 4°C until sequencing, to preserve high molecular weight for long-read sequencing.

DNA and RNA were quantified with the Qubit dsDNA Broad Range and RNA High Sensitivity assays respectively (Thermo Fisher Scientific), and their integrity assessed on a TapeStation system using Genomic DNA and RNA ScreenTape (Agilent Technologies).

### 4. Long-read ONT sequencing

Details of all DNA and RNA sequencing runs are provided in Table S1.

#### 4.1. DNA sequencing

For ONT DNA sequencing runs, the three biological replicates of each tissue were sequenced for the highly infected level, whereas a single sample per tissue was sequenced for the lightly and moderately infected levels (n = 20 samples total: 3 replicates × 4 tissues for the highly infected level, plus 1 replicate × 4 tissues for each of the lightly and moderately infected levels). Samples from the not infected control tank were not included in DNA sequencing. This design was intended to allow intra-host viral variant calling analysis where viral load was sufficient, while limiting sequencing effort at lower infection levels.

Libraries were prepared using the Ligation Sequencing Kit with Native Barcoding 24 V14 (SQK-NBD114.24) and loaded onto R10.4.1 flowcells. Five sequencing runs were performed, each multiplexing four samples on the same flowcell (Table S1). For the first three runs, adaptive sampling was enabled to enrich OsHV-1 reads in real time against the OsHV-1 reference genome (accession: GCA_964656485.1) (Table S1).

#### 4.2 RNA sequencing

For RNA sequencing, the three biological replicates of each tissue were sequenced for all four infection levels, including the samples from the not infected control tank, giving a total of 48 samples across 23 sequencing runs (Table S1). Because OsHV-1 DNA was not present in not infected oysters, this group was not included in DNA sequencing. RNA sequencing was nonetheless performed on these samples to provide a host transcriptomic baseline for the unexposed condition.

The first two libraries (run 1, replicate 1 of highly infected gills; run 2, replicate 1 of moderately infected gills) were prepared with the cDNA-PCR Sequencing Kit (SQK-PCS114) and sequenced individually. The remaining 21 libraries were prepared with the cDNA-PCR Sequencing Kit with Barcoding (SQK-PCB114.24), multiplexing three samples per flow cell. All libraries were loaded onto R10.4.1 flow cells. Three flow cells were washed and reloaded to increase output (runs 3 and 4, replicates 2 and 3 of highly and moderately infected gills; run 9, replicates 1 to 3 of highly infected muscle) (Table S1).

Sequencing was performed on MinION Mk1B or Mk1C devices controlled by MinKNOW v24.06.14.

### 5. Data analysis

Raw signal data (POD5 format) from all sequencing runs were transferred to Datarmor, the Ifremer high-performance computing cluster, for downstream analysis.

#### 5.1. OsHV-1 genomic characterization

##### Basecalling and reads filtering

Raw POD5 files from each DNA sequencing run were basecalled with Dorado v0.9.1 (Oxford Nanopore Technologies) using the dna_r10.4.1_e8.2_400bps_sup@v5.0.0 model, with -- batchsize 256, --trim all to remove adapters and barcodes, and --kit-name SQK-NBD114-24 to perform barcode classification in-line during basecalling. The resulting classified BAM files were demultiplexed with dorado demux --no-classify, which assigns reads to their barcode based on the classification already recorded during basecalling, and converted to FASTQ using the -- emit-fastq option. Read quality control and removal of residual adapter and barcode sequences were performed with fastplong v0.2.2 (Chen, 2023), using default quality and length filtering thresholds (minimum base quality Q15, reads with >40% unqualified bases discarded, minimum read length 20 bp), specifying the start adapter (-s CCTGTACTTCGTTCAGTTACGTATTGCT) and the end adapter (-e ACGTAACTGAACGAAGTACAGG), together with a FASTA file containing all barcode sequences used (--adapter_fasta).

##### Read sorting and classification

To assign each genomic read to its organism of origin (i.e. *M. gigas* or OsHV-1), cleaned reads were aligned simultaneously against the *M. gigas* nuclear and mitochondrial genome (GenBank: GCF_963853765.1 and NC_001276 respectively) and the OsHV-1 genome using Minimap2 version 2.30 (Li, 2018) with -ax map-ont, --sam-hit-only and --secondary=no. The OsHV-1 reference (GenBank GCA_964656485.1) was assembled in a previous study from an isolate collected at the same site (La Floride) in 2020 (Dotto-Maurel, Pelletier, et al., 2025). SAM files were converted, sorted and indexed BAMs using Samtools version 1.22.1 (Danecek et al., 2021). Custom scripts then partitioned the reads into four categories based on the presence or absence of an alignment record against each reference. These comprised reads that aligned uniquely to either *M. gigas* or OsHV-1, reads that aligned to both references (likely chimeric reads arising from ligation during library preparation), and reads that aligned to neither. The latter were further characterized as detailed in Supplementary File 1.

Because adaptive sampling was applied to part of sequencing runs (Table S1), a substantial fraction of the rejected, non-viral reads were shorter than 500 bp and were poorly handled by the alignment parameters above. These unaligned short reads were therefore realigned with -ax sr -- sam-hit-only --secondary=no. Residual oyster reads may also have escaped detection because of divergence from the reference or incompleteness of the assembly; short-read alignments were thus additionally performed against all *M. gigas* genome assemblies publicly available in NCBI (GCA_011032805.1, GCA_025765685.3, GCA_963853805.1, GCA_005518195.2, GCA_000297895.2, GCA_902806645.1). SAM files were processed as described above and the resulting FASTQ files were merged with those obtained from the initial alignment. Read counts, base counts and N50 values were computed with SeqKit v2.8.1 (Shen et al., 2024).

##### OsHV-1 genome de novo assembly

To generate a high-quality OsHV-1 genome assembly, non-*M. gigas* reads were pooled across all the different sequenced libraries and assembled with Canu version 2.3 (Koren et al., 2017) with the parameter genomeSize set to 0.204 m. A custom script was then used to reconstruct the canonical OsHV-1 genome architecture in two steps. First, the known OsHV-1 regions of the reference genome (GenBank: GCA_964656485.1) (Dotto-Maurel, Pelletier, et al., 2025) were aligned against the assembled contigs with BlastN v2.12.0 (Altschul, 2014) and the resulting hits exported in GFF format. Secondly, the GFF and the contig Fasta files were parsed with BEDtools v2.30.0 (Quinlan & Hall, 2010), Seqtk version 1.4 (https://github.com/lh3/seqtk) and BlastN to identify the viral regions, validate them by length and reorder them into the canonical OsHV-1structure (TR_L_ - U_L_ - IR_L_ - X - IR_S_ - U_S_ - TR_S_).

The assembly was polished with Medaka v2.2.1 (medaka_consensus, model r1041_e82_400bps_sup_v5.0.0) using the same set of non-*M. gigas* reads. Assembly accuracy was then assessed by realigning all non-*M. gigas* reads to the polished genome with Minimap2 v2.30 and processing the alignments with SAMtools v1.22.1.

##### Phylogenetic placement

To place this OsHV-1 genome within the broader OsHV-1 phylogeny, the polished assembly was aligned with all publicly available OsHV-1 genomes retrieved from NCBI (https://www.ncbi.nlm.nih.gov/datasets/genome/) and the MopSeq database (Battistel et al., 2025) (https://mopseq-db.ifremer.fr/table_pat/Viruses/Ostreid%20herpesvirus-1/) using MAFFT v7.520 with the --auto option, which automatically selects the most appropriate alignment strategy based on sequence number and length. A maximum likelihood tree was then reconstructed with IQ-TREE v2.0.3 (Minh et al., 2020) using -m MFP -B 1000, and branch support was assessed by 1000 ultrafast bootstrap replicates. ModelFinder (Kalyaanamoorthy et al., 2017) selected HKY+F+R7 as the best-fit substitution model according to the Bayesian information criterion. The tree was midpoint-rooted and visualized with FigTree v1.4.4 (https://tree.bio.ed.ac.uk/software/figtree/).

##### OsHV-1 genome annotation

This new OsHV-1 genome was then annotated using pairwise alignment with a previously annotated OsHV-1 genome (Dotto-Maurel, Luyer, et al., 2025) with Geneious version 2025.0.2 (https://www.geneious.com) using the transfer annotation option. The generated GFF file was then converted to GTF format for subsequent analysis using the agat_convert_sp_gff2gtf.pl script from AGAT version 1.4.0 (Dainat, 2024).

##### OsHV-1 isomers identification

The four major isomers (P: U_L_→/ U_S_→; I_S_: U_L_→/ U_S_←; I_LS_: U_L_←/ U_S_ ← and I_L_: U_L_ ←/ U_S_→) were reconstructed from the new OsHV-1 assembly by permuting the orientation of the U_L_ and U_S_ regions. For each configuration, the terminal 40 kb of the linear genome (encompassing the U_L_-IR_L_-X-IR_S_-U_S_ region) was extracted. Supporting reads were identified by first discarding non-*M. gigas* reads shorter than 9 kb, then aligning the remaining reads with Minimap2 using - ax map-ont, --sam-hit-only and --secondary=no parameters, against a single reference combining the four isomer sequences. Only alignments with MAPQ > 40 and spanning part of U_L_ and U_S_ and the full IR_L_-X-IR_S_ region were retained using SAMtools and Geneious.

##### OsHV-1 genomic intra-variant calling

To assess whether viral genotypes differed among tissues, variant calling was performed with Medaka v2.2.1 on OsHV-1 reads from highly infected oysters against the newly assembled OsHV-1 reference. VCF files were then analyzed in RStudio version 2024.09.1 (https://posit.co/download/rstudio-desktop/). For each variant, the variant allele frequency (VAF) was calculated from the VCF SR field (number of reads best supporting each allele, by strand) and the DPSP field (total read depth spanning the position ± 25 bp, including ambiguous reads):

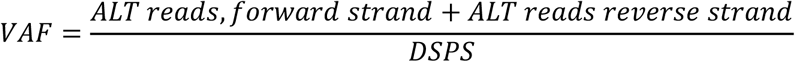

Only variants reaching a minimum coverage depth (DPSP) of 5× were retained for analysis, to limit calling noise associated with low-coverage positions. Single-nucleotide substitution variants (SNVs) were classified as transitions (A↔G, C↔T) or transversions (all other substitutions); insertions/deletions were excluded from this classification.

The relative effects of host individual (oyster) and tissue on the structure of viral variants were tested using a permutational multivariate analysis of variance (PERMANOVA) on a Bray-Curtis dissimilarity matrix calculated from the VAF profiles of each sample, with oyster and tissue included as explanatory factors and tested as marginal effects (Anderson, 2001). Because only three oysters were available and tissues sampled within the same individual are not statistically independent, the resulting p-values were interpreted as indicative rather than exact, and the relative effect sizes (R²) were used as the primary basis for comparing the two factors. As a descriptive complement, pairwise similarity of variant profiles was also summarized using Jaccard dissimilarity on variant presence/absence, contrasting sample pairs from the same oyster with sample pairs from the same tissue.

#### 5.2. Long-read transcriptomic basecalling, reads quality and sorting

Raw pod5 files from the cDNA sequencing runs were basecalled with Dorado using the same DNA model as for the DNA runs (dna_r10.4.1_e8.2_400bps_sup@v5.0.0), with the --no-trim option to preserve primer sequences required by Pychopper for full-length read identification and orientation. Full-length cDNA reads were then identified, oriented and trimmed with Pychopper v2.7.10 (https://github.com/epi2me-labs/pychopper) with the default parameters across all 48 libraries. Quality control and removal of residual adapters were performed with Fastplong specifying the start adapter (--start_adapter CTTGCGGGCGGCGGACTCTCCTCTGAAGATAGAGCGACAGGCAAG) and a FASTA file listing all adapter and barcode sequences (--adapter_fasta) (https://nanoporetech.com/document/chemistry-technical-document#barcode-sequences).

To determine the organism of origin of each read, the cleaned cDNA reads (i.e., reads cleaned with Pychopper and Fastplong as described above) were aligned to the *M. gigas* nuclear and mitochondrial reference genomes (GenBank: GCF_963853765.1 and NC_001276, respectively), and to the newly assembled OsHV-1 genome, with Minimap2 using the parameters -ax splice - k14 -G10k -ub --sam-hit-only --secondary=no. Alignments were converted, sorted, and indexed using SAMtools. Custom scripts were then used to partition reads assigned to *M. gigas*, to OsHV-1, and to neither reference. Unassigned reads were subsequently screened against all publicly available *M. gigas* assemblies (accessions as above) using the same alignment strategy to recover any remaining oyster transcripts.

#### 5.3. OsHV-1 long-read transcriptomic analysis

##### OsHV-1 transcript identification

To reconstruct the viral transcriptome, OsHV-1 cDNA raw basecalled reads were aligned to the non-redundant assembly (P isomer, lacking TRL and TRS regions) with Minimap2, using the parameters recommended for LoRTIA (-ax splice -Y -C5 --cs). LoRTIA v0.9.9 (https://github.com/zsolt-balazs/LoRTIA) was chosen because it resolves transcription start and end site (TSS/TES) boundaries even in regions of overlapping transcription units, a situation typical of compact, intron-poor viral genomes, and has previously been applied to other herpesvirus transcriptomes (Tombácz et al., 2019; Torma et al., 2023). Although OsHV-1 transcripts are predominantly intronless, the splice-aware mode was retained because it improves the handling of transcript termini through soft clipping and provides the base-level alignment information required for accurate detection of TSS and TES. Alignments were thus processed with LoRTIA (-5 TGCCATTAGGCCGGG --five_score 14 --check_in_soft 15 -3 AAAAAAAAAAAAAAA --three_score 14 -s poisson -f true). A transcriptome FASTA file was generated from the LoRTIA GFF output, and transcripts whose start and end positions differed by less than 150 bp were merged, retaining the longest isoform.

##### OsHV-1 transcript classification

To characterize the organization of the OsHV-1 transcriptome, a transcript-to-gene correspondence table was first generated. For each transcript, the percentage of overlap with every annotated gene was computed based on strand information derived directly from the LoRTIA alignment, with overlaps on the antisense strand recorded as negative values.

Each transcript was classified according to a decision tree based on the number of annotated genes it overlapped, the extent of that overlap (using a 90% threshold), and the relative orientation of transcript and gene(s) (Figure 9.A). Transcripts overlapping no gene were termed intergenic. Transcripts overlapping a single gene by more than 90%, in the same orientation, were classified as monocistronic; those overlapping a single gene by more than 90% in the opposite orientation were classified as antisense. Transcripts overlapping several genes, all by more than 90% and in the same orientation, were classified as polycistronic; when genes with discordant orientations were involved, transcripts were classified as complex. Transcripts falling below the 90% overlap threshold, whether overlapping one or several genes, were assigned to an overlapping category and further subdivided based on the number of genes involved, the overlap achieved for each gene, and their relative orientation, yielding partial monocistronic, partial antisense, partial bicistronic, mixed monocistronic, complex partial, complex mixed monocistronic and complex mixed polycistronic subtypes (Figure 9.B). Major categories (monocistronic, polycistronic, antisense, complex, intergenic) follow the terminology established in a previous study (Boldogkői, Moldován, et al., 2019); the decision tree and overlap thresholds applied to define them, as well as the overlapping subtypes, were established for this study.

**Figure 9:**
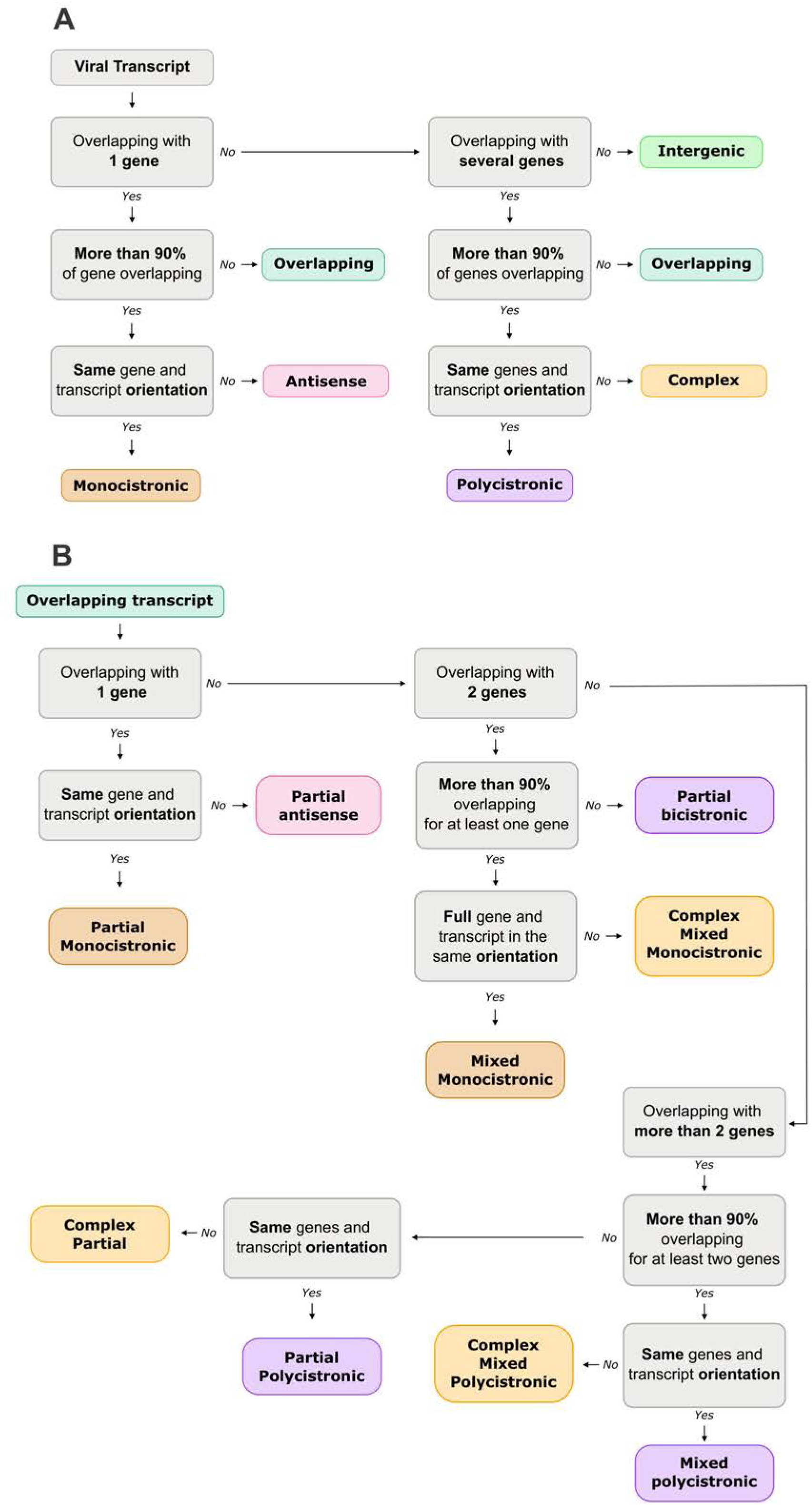
Decision tree for classifying OsHV-1 viral transcripts based on gene overlap. Grey boxes are binary decision steps (Yes/No); colored boxes are final transcript categories. A. First-level classification. Each transcript is first tested for overlap with one or several genes (or none, → Intergenic). For single-gene overlaps, transcripts with ≤90% overlap are classified as Overlapping; those with >90% overlap are further split by gene/transcript orientation into Antisense (opposite strand) or Monocistronic (same strand). The same logic applies to multi-gene overlaps, yielding Complex (mixed orientation) or Polycistronic (same orientation, further denoted bicistronic, tricistronic, etc. according to the number of genes involved) for transcripts overlapping each gene by more than 90%. B. Sub-classification of overlapping transcripts. Transcripts assigned to Overlapping in (A) are further split by number of genes overlapped, percentage overlap per gene, and relative orientation, yielding partial, mixed, and complex mixed subtypes; no transcripts were assigned to the partial antisense category.

##### OsHV-1 transcript quantification

To quantify viral transcript abundance, OsHV-1 cDNA reads from each sample were aligned to the transcriptome produced by LoRTIA with Minimap2 using the parameters -ax map-ont, -p 0 and -N 10, so that multiple valid alignments per read were retained. A custom script parsed CIGAR strings to compute reference coverage for each alignment and reassigned the 0×100 flag so that only the best alignment of each read was retained as primary. Primary alignments were then quantified with NanoCount v.1.1.0post2 (Gleeson et al., 2022),, using the “primary” alignment field to define the best alignment for each read (rather than the default alignment score) and disabling the default minimum alignment length and minimum query fraction aligned filters (both set to 0), since alignment selection had already been performed by the custom script described above.

##### Count matrix normalisation

Viral and (see below) host transcript count matrices were merged. Infection level (not infected, lightly, moderately and highly infected) and tissue (gills, mantle, digestive gland, muscle) were encoded as ordered factors and combined into a single composite factor, and raw counts were normalized with DESeq2 v1.46.0 (Love et al., 2014) under a ∼ group design. A variance-stabilizing transformation (vst) was then applied for downstream analyses. Because viral transcripts represented at most 20.5% of sequenced bases in the most highly infected samples, size factors estimated from the merged host-viral count matrix were compared to those estimated from host transcripts alone (Pearson’s r = 1.00). The near-perfect agreement between the two indicated that joint normalization did not introduce a detectable composition bias, and normalization was therefore performed on the merged matrix throughout.

##### OsHV-1 ORF expression kinetics

To assess OsHV-1 RNA coverage, Minimap2 alignments to the non-redundant OsHV-1 assembly were converted into per-base coverage with “bedtools genomecov” using BEDtools. Coverage values were expressed as coverage per billion sequenced bases, calculated by dividing the coverage at each position by the total number of bases sequenced in the library and multiplying by 10⁹.

To quantify gene expression at the ORF level across tissues and infection levels, the 48 cDNA libraries were aligned to the *M. gigas* nuclear and mitochondrial genomes and to the newly assembled non-redundant OsHV-1 assembly with Minimap2 using parameters -ax splice -k14 - G10k -ub --sam-hit-only --secondary=no. Reads were assigned to features with featureCounts v2.0.8 (Y. Liao et al., 2014) using parameters -L -g gene -t exon --minOverlap 150 -O, so that reads spanning several genes were counted for every overlapping feature and transcripts covering more than one ORF were not undercounted. This ORF-level count matrix is independent of the transcript-level count matrix; it was normalized with DESeq2 and vst-transformed following the procedure described above, with the same composite group factor. For each of the annotated OsHV-1 ORFs, a stage-level expression profile was obtained by taking the median vst value across all tissues and replicates within each infection level (not infected, lightly, moderately, highly infected). These four-point profiles were clustered using the k-means method (k = 3, nstart = 50), which is consistent with the immediate-early, early and late kinetic classes that are usually found in herpesvirus lytic transcription programs (Dotto-Maurel, Arzul, et al., 2025; Dotto-Maurel, Le Luyer, et al., 2025).

##### Composition of the viral transcriptome

The relative contribution of tissue and infection level to the composition of the viral transcriptome was assessed by redundancy analysis (RDA). Because infection level is defined from viral load, an ordination based on absolute transcript abundance would trivially separate libraries by infection level regardless of any genuine change in transcriptome composition. Libraries from not infected oysters were therefore excluded from this analysis, as viral counts of zero throughout preclude a compositional transformation. Prior to RDA, DESeq2-normalized counts of the OsHV-1 transcripts across the remaining libraries were converted to row-wise proportions of the total viral counts in each library, and Hellinger-transformed (Legendre & Gallagher, 2001), thereby isolating shifts in the relative composition of the viral transcriptome from variation in overall viral abundance across libraries.

Because the four tissues sampled from a given oyster are not statistically independent, whereas oysters themselves are, two different permutation schemes were used to test each factor with anova.cca (by = “margin”, 9999 permutations). The effect of infection level, which varies between oysters, was tested by permuting entire oysters (keeping their four tissues together); the effect of tissue, which varies within an oyster, was tested by permuting tissues only within each oyster.

##### OsHV-1 transcript detection across tissues

To assess whether OsHV-1 transcripts were detected across all four tissues, a transcript was considered detected in a given tissue if DESeq2-normalized counts reached ≥5 in at least 2 of the 9 libraries from that tissue, using libraries from infected oysters only (lightly, moderately and highly infected; not infected libraries were excluded as viral counts were uniformly zero).

To test whether incomplete detection across tissues reflected tissue-specific expression or simply differences in viral sequencing depth between libraries, raw counts were equalized to the lowest viral read depth among libraries with ≥50 viral reads by random subsampling (multinomial rarefaction), and libraries with fewer than 50 viral reads were excluded from this analysis. The detection criterion above was reapplied to the rarefied counts, and the observed proportion of transcripts with incomplete detection was compared to a null distribution generated by randomly permuting tissue labels across libraries while keeping rarefied count values unchanged (200 permutations).

The association between detection completeness (detected in all four tissues vs. not) and transcript category was tested with a χ² test. The relationship between transcript length and detection completeness was assessed by logistic regression, with transcript category included as a covariate to test whether the two factors contributed independently; nested models were compared by likelihood ratio tests.

##### Comparison of OsHV-1 polycistronic and monocistronic transcript abundance

For ORFs contained in both a polycistronic and a monocistronic transcript, DESeq2-normalized counts were averaged across replicates for each transcript, tissue and infection level. Abundance was compared between the two transcript types across matched ORFs using a paired Wilcoxon signed-rank test. The change in abundance between moderately and highly infected oysters, the two infection levels at which both transcript types were consistently detected, was compared between monocistronic and polycistronic transcripts using a paired Wilcoxon signed-rank test on per-ORF fold-changes.

##### OsHV-1 transcript differential expression between tissues

Differential expression between tissues was assessed separately for each infection level using DESeq2, with size factors estimated from the combined host and viral count matrix (design: ∼ Tissue + Infection level + Tissue:Infection level). For each pair of tissues within a given infection level, a numeric contrast vector was constructed from the relevant model coefficients, explicitly accounting for which tissue served as the reference level in the design, so that no coefficient was included or omitted inconsistently between the two tissues being compared. To avoid inflating the false discovery rate through post-hoc filtering on unshrunken effect sizes, log2 fold changes were shrunk using the ashr method (lfcShrink) (Stephens, 2017) prior to filtering. Viral transcripts with a shrunken absolute log2 fold change above 2 and an adjusted p-value below 0.05 were considered differentially expressed.

5.4. *M. gigas* long-read transcriptomic analysis

##### M. gigas transcript quantification

Unlike OsHV-1, oyster transcripts are extensively spliced, so that reads must be assigned to isoforms sharing overlapping exons. FLAIR v.2.2.0 (Tang et al., 2020) was therefore used, as it performs splice-aware quantification and relies on known splice junctions to disambiguate isoform assignment. Cleaned *M. gigas* cDNA reads (i.e., reads assigned to *M. gigas* as described in section 5.2) were quantified with the “quantify” command and default parameters against the *M. gigas* reference transcriptome (accession number: GCF_963853765.1). The resulting count matrix was merged with the viral one and normalized as described in section 5.2.

##### Effects of tissue and infection level on the host transcriptome

The relative contribution of tissue and infection level to the host transcriptome was assessed by distance-based redundancy analysis (db-RDA). A principal coordinate analysis was first performed on the host vst matrix using Euclidean distance, and the first six axes were retained as the response matrix in a db-RDA with tissue and infection level as explanatory variables. The two factors were tested with anova.cca (by = “margin”, 9999 permutations) using the same permutation schemes as described in section 5.2 (permutation of entire oysters for infection level; permutation of tissues within each oyster for tissue).

##### Weighted gene co-expression network analysis

Oyster transcripts were extracted from the vst matrix and those with a variance below 0.15 across samples were discarded (17,824 of 46,043 transcripts retained), to focus the co-expression analysis on transcripts with sufficient variability to inform module detection. A weighted gene co-expression network analysis (WGCNA) was performed on the remaining transcripts across all libraries with the WGCNA package v1.73 (Langfelder & Horvath, 2008). Sample traits comprised tissue, infection level and viral load, the latter expressed as OsHV-1 genome copies per ng of total DNA measured by qPCR. A soft-thresholding power of 10 was selected according to the scale-free topology criterion (model fit 0.85, mean connectivity 201), and modules were defined with cutreeDynamic using a minimum module size of 50 transcripts. Module membership was computed for each module, and correlations between module eigengenes and sample traits were assessed using Spearman correlation as an initial screening step. Modules with an eigengene correlated with infection level and/or viral load (p < 0.01, r² > 0.5) were retained for further analysis. The significance of these associations, and of the tissue × infection level interaction for each retained module, was then assessed using linear mixed models on module eigengenes with oyster as a random effect, to account for the non-independence of the four tissues sampled from the same individual.

##### Functional enrichment of WGCNA modules

Functional enrichment was performed on the genes of the retained modules using the gene ontology (GO) annotation previously assigned to the *M. gigas* genome (Dotto-Maurel, Le Luyer, et al., 2025), with a rank-based approach and adaptive clustering implemented in GO_MWU (Wright et al., 2015), applying a Mann-Whitney U test to each module against the go.obo database (January 2025 release). Terms with an adjusted p-value below 0.01 were retained, and enrichment was summarized by the negative logarithm of the adjusted p-value and the proportion of significant genes per term.

##### Host differential expression

Host differential expression was assessed with DESeq2. To compare the response to infection within each tissue, each infection level was compared with the not infected control separately for each of the four tissues. To identify tissue-specific features of the host response at the highly infected stage, the four tissues were compared pairwise using a design restricted to highly infected libraries and accounting for individual oysters (design: ∼ oyster + tissue). Transcripts differentially expressed between a given tissue and each of the three other tissues, in the same direction, were considered specific to that tissue. In both analyses, log2 fold changes were shrunk using the ashr method (lfcShrink) (Stephens, 2017) prior to filtering, and transcripts with a shrunken absolute log2 fold change above 2 and an adjusted p-value below 0.05 were considered differentially expressed. The resulting sets of differentially expressed transcripts were compared with the WGCNA modules correlated with infection level and viral load (set to 0 for not infected samples).

## DATA AVAILABILITY

The raw Oxford Nanopore sequencing data (gDNA and cDNA reads) generated in this study will be deposited in the Ifremer data repository as soon as possible and will be made publicly available in the European Nucleotide Archive (ENA) upon acceptance. The newly assembled OsHV-1 genome will be deposited in GenBank, and its accession number will be provided upon acceptance. All scripts used in this study are available at https://gitlab.ifremer.fr/asim/in_vivo_lytic_phase_oshv-1_long_reads.

## ACKNOWLEDGEMENTS

We thank the staff of the Ifremer ASIM laboratory in La Tremblade for their help with sampling. We are grateful to Frédéric Girardin and his team at the Plateforme des Mollusques Marins de La Tremblade (PMMLT), and to Virginie François and her team at the Plateforme des Mollusques Marins de Bouin (PMMB), for producing the oysters used in this study. We also thank the SEBIMER team for maintaining the bioinformatics tools, and all members of the ASIM laboratory for their support. We acknowledge the Pôle de Calcul et de Données Marines (PCDM; https://wwz.ifremer.fr/en/Research-Technology/Research-Infrastructures/Digital-infrastructures/Computation-Centre) for providing DATARMOR computing and storage resources. A.D.-M. was supported by a grant from the Ifremer Scientific Board. The funders had no role in study design, data collection and interpretation, or the decision to submit the work for publication.

---

**Figure S1:**
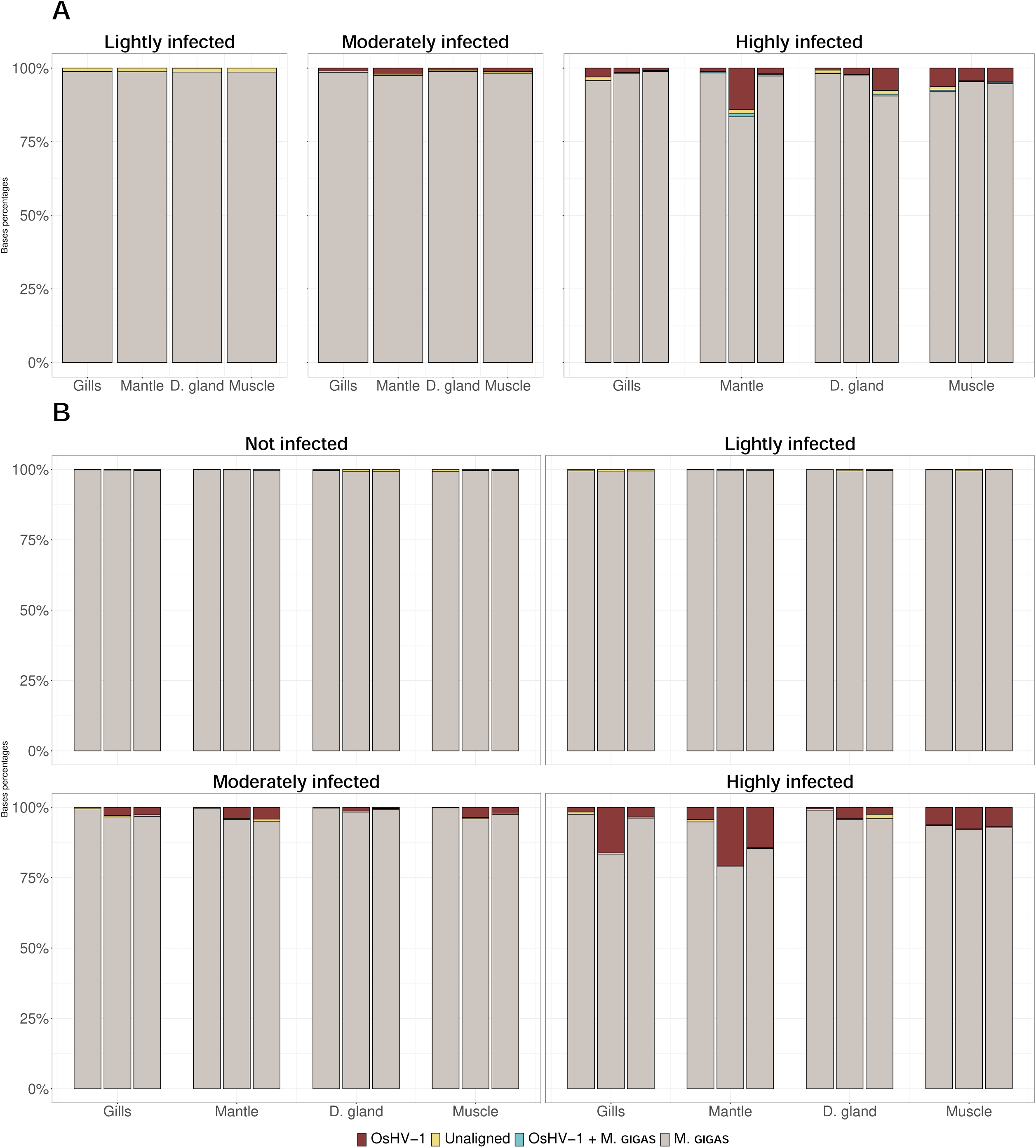
Classification of sequenced bases for gDNA (A) and cDNA (B) sequencing runs. The bar charts show the proportion of sequenced bases according to their alignment in four oyster tissues (gills, digestive gland, mantle and muscle). Each bar represents a sequencing library corresponding to one sample per tissue and condition for genomic DNA (gDNA) and three biological replicates per tissue and condition for complementary DNA (cDNA). Bases mapping exclusively to the *M. gigas* genome are shown in light grey, bases mapping to both OsHV-1 and *M. gigas* in blue, unaligned bases (those not mapping to either genome) in yellow, or bases mapping exclusively to the OsHV-1 genome in maroon.

**Figure S2:**
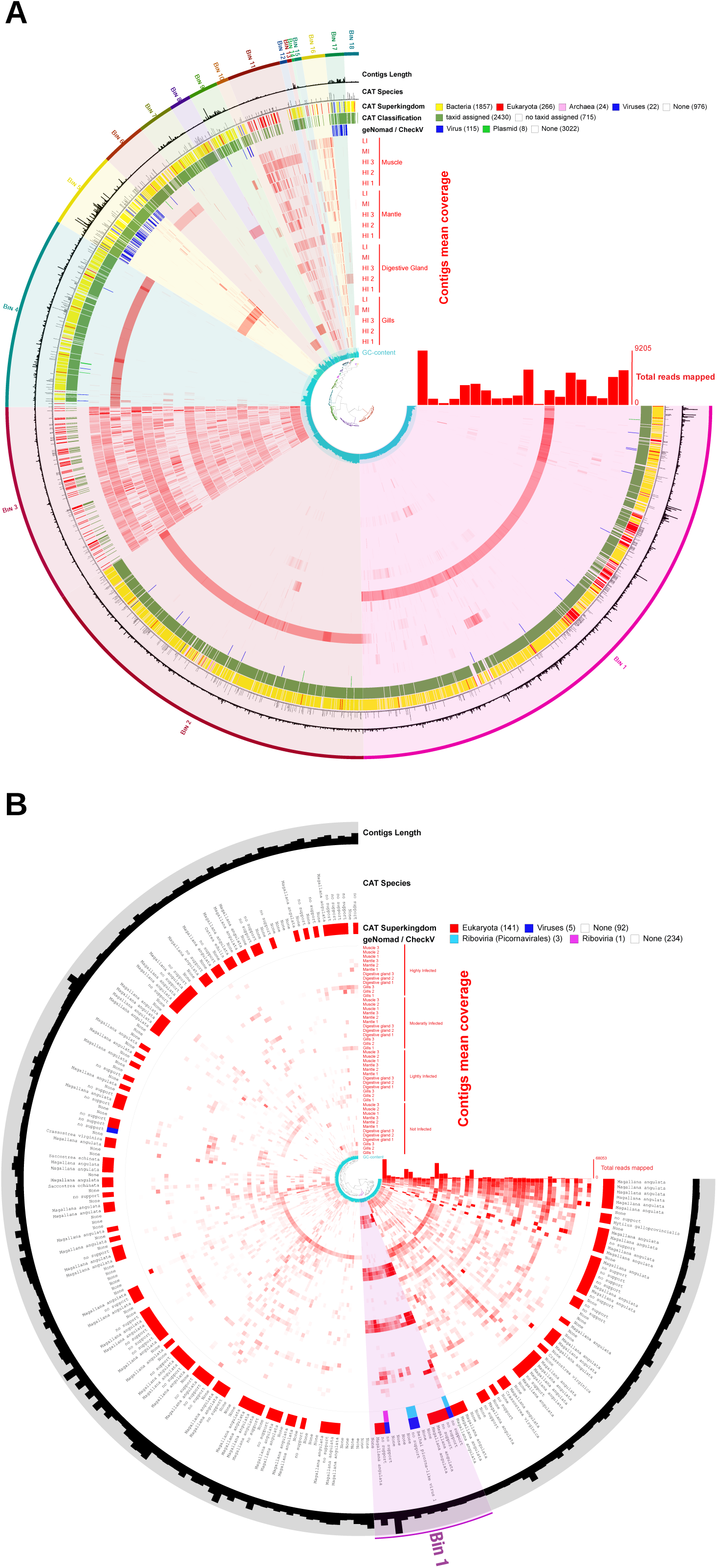
Composition, taxonomy and coverage of contigs assembled from unaligned reads. Each radial item represents one contig, ordered by hierarchical clustering based on sequence composition and differential coverage; bins delineated from this clustering are indicated in the outer ring. In both panels, from the outside inwards: contig length; CAT taxonomic assignment at the species and superkingdom levels; classification of contigs as viral or plasmid sequences by geNomad and CheckV; mean coverage of each contig in each library; and GC content. The central dendrogram shows the clustering of contigs, and the bar plot gives the total number of reads mapped per library. Numbers in the legend indicate the number of contigs in each category. A. Genomic contigs (n = 3,145 of at least 1 kb), grouped into 18 bins, with their mean coverage in the 20 genomic libraries corresponding to four tissues (muscle, mantle, digestive gland and gills) and three infection levels (LI, lightly infected; MI, moderately infected; HI 1 to HI 3, three biological replicates of highly infected oysters); maximum number of reads mapped per library, 9,205. A fourth ring indicates whether a taxon identifier was assigned by CAT. B. Transcript contigs (n = 238, out of 1,301 assembled), grouped into a single bin, with their mean coverage in the 48 cDNA libraries corresponding to four infection levels (not infected, lightly, moderately and highly infected), four tissues (muscle, mantle, digestive gland and gills) and three biological replicates; maximum number of reads mapped per library, 68,053.

**Figure S3:**
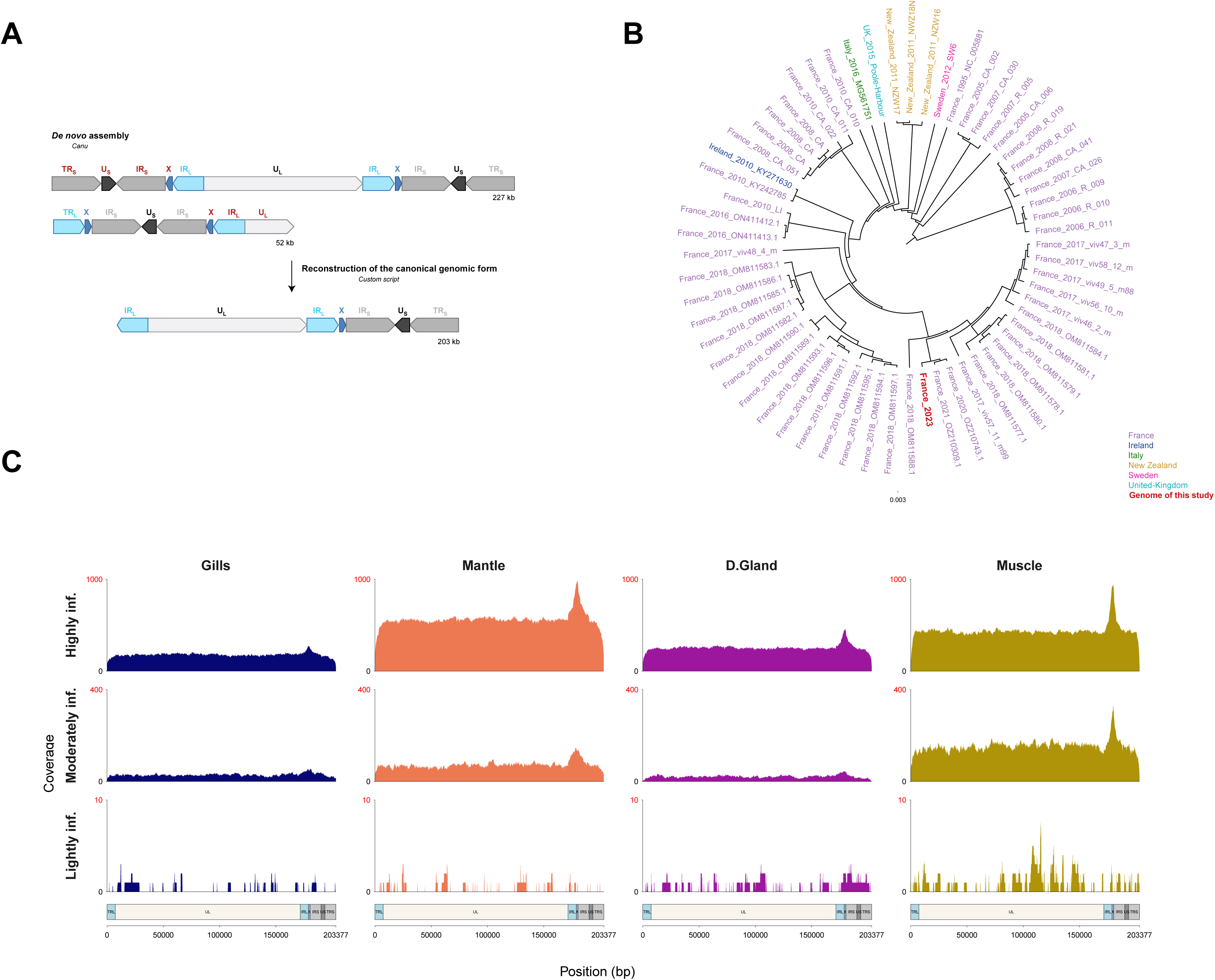
*De novo* assembly, phylogeny and reads mapping. A. Schematic representation of *de novo* assembly contigs and reconstruction of the final genome. Names of redundant regions were colored in darkred. B. Phylogenetic tree of all publicly available, non-redundant OsHV-1 genomes. All non-redundant OsHV-1 genomes isolated from *M. gigas* and available in public databases were aligned with the polished genome assembled in this study to construct the phylogenetic tree. Genome labels include the country and year of isolation, as well as the GenBank accession number or assembly name if not yet released. The genome generated in this study is highlighted in red and bold. C. Read coverage across the assembled and polished OsHV-1 genome. Each column corresponds to one of four tissues and each row shows a different infection level: highly infected (top), moderately infected (middle) or lightly infected (bottom). For the highly infected condition, the average read coverage across the three oysters for each tissue is shown. The y-axis shows the read depth in base pairs, scaled from 0 to 2,500 for high infection, 0 to 400 for moderate infection, and 0 to 10 for low infection. The x-axis shows the OsHV-1 genome position in base pairs. Beneath each column, colored bars denote genome regions: U_L_ (beige), TR_L_/IR_L_ (light blue), X (dark blue), TR_S_/IR_S_ (light grey) and U_S_ (dark grey).

**Figure S4:**
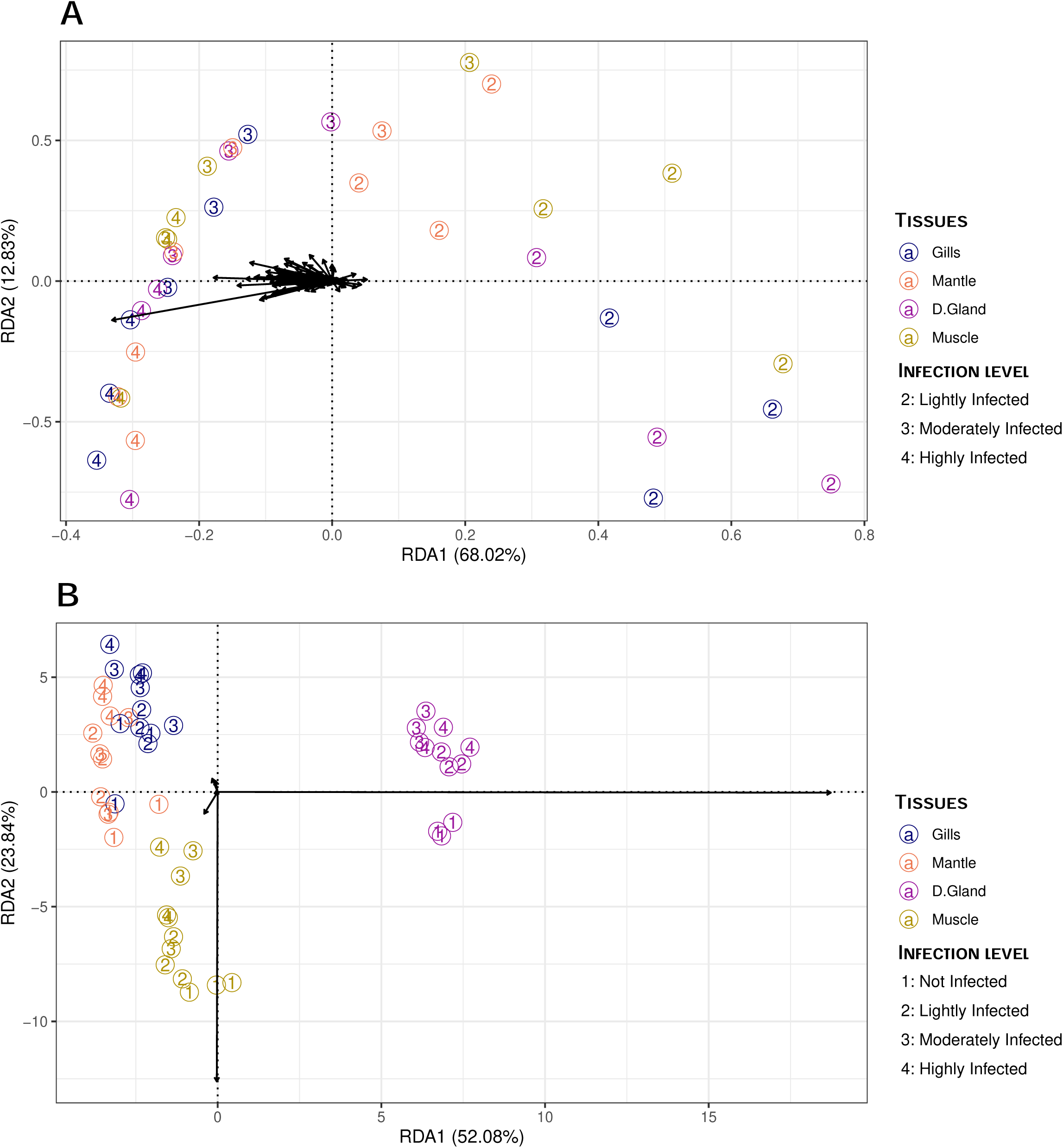
Redundancy Analysis (RDA) plot. Each point represents a library, with numbers indicating infection levels: 1 = not infected; 2 = lightly infected; 3 = moderately infected; and 4 = highly infected. The colors of the dots indicate tissue types: dark blue for gills, orange for the mantle, purple for the digestive gland, and dark yellow for muscle. A. RDA using viral gene expression only. RDA1 explains 68.02% of the variance and RDA2 explains 12.83% of the variance. B. RDA using host gene expression only. The x-axis represents the first RDA component (RDA1), explaining 52.08% of the variance; the y-axis shows the second component (RDA2), accounting for 23.84% of the variance.

**Figure S5:**
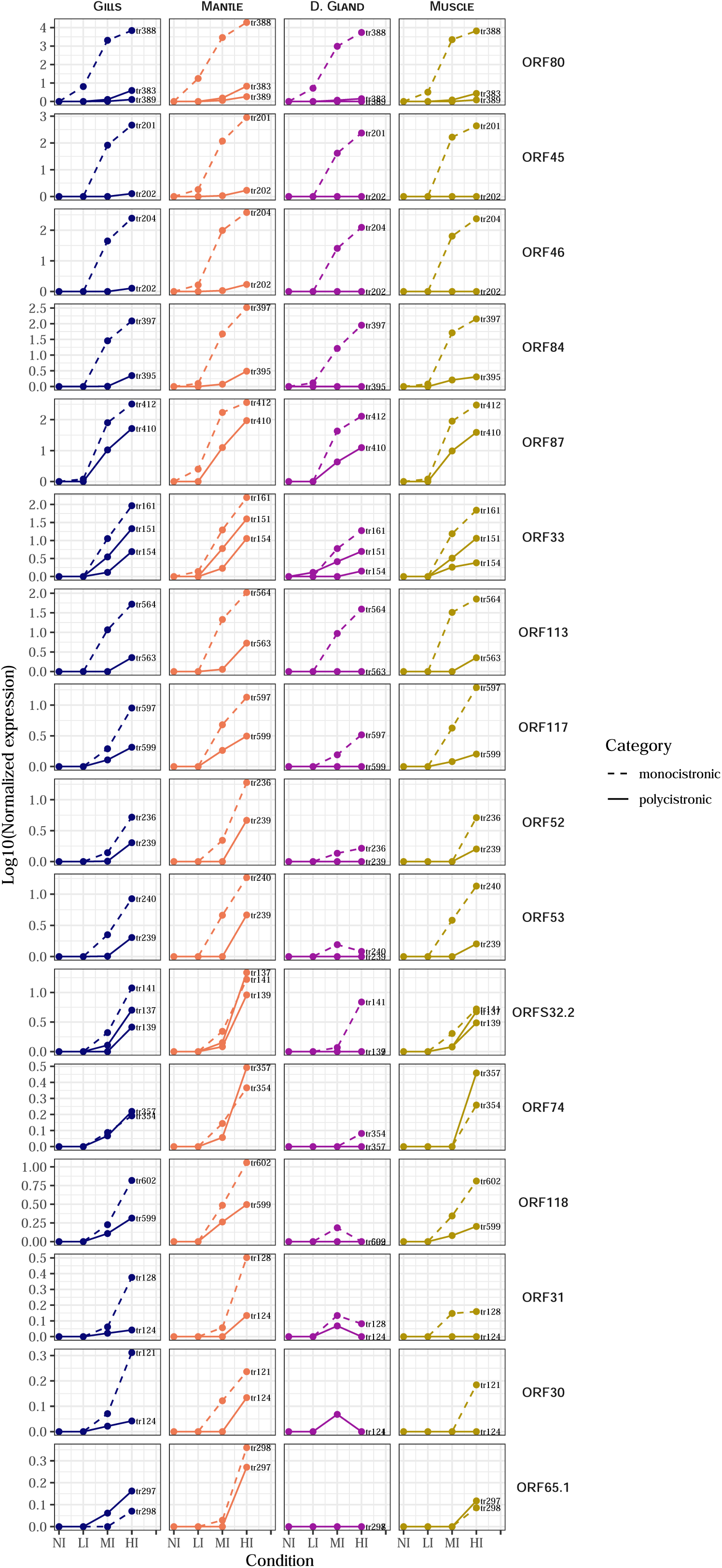
Expression of monocistronic and polycistronic transcripts. Each row panel represents a different gene ordered by expression level, while each column corresponds to a specific tissue. The Y-axis of each facet shows the log10 of normalized expression, and the X-axis indicates the infection condition. Expression from monocistronic transcripts is represented by dashed lines, whereas expression from polycistronic transcripts is shown with solid lines. Each point and line is color-coded according to the corresponding tissue.

**Figure S6:**
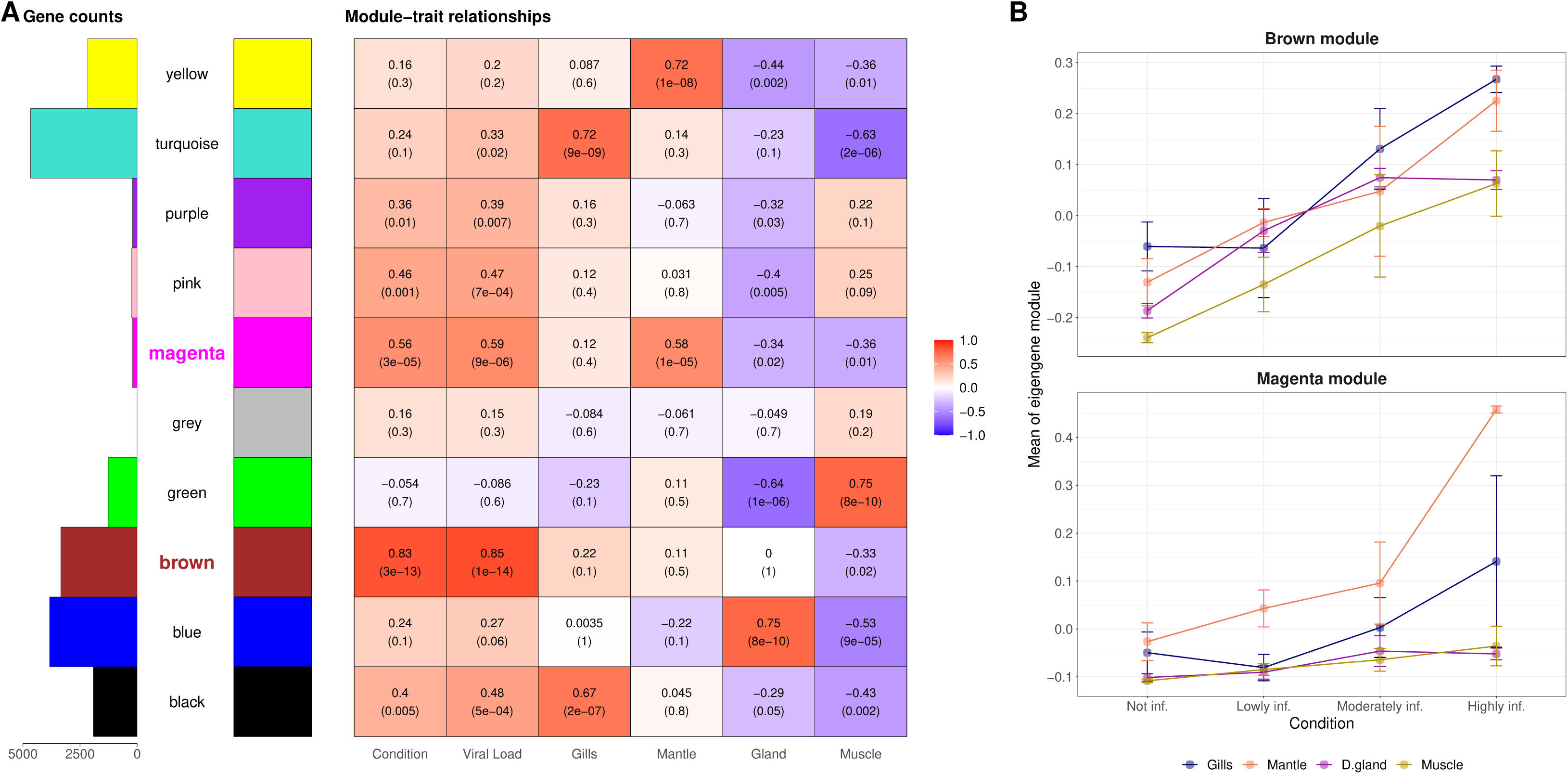
WGCNA modules description, statistical analysis of oyster gene expression and mean expression of infection-specific modules. A. The left panel shows the number of genes per module, with the X-axis indicating gene count, while the right panel represents the correlation between modules and traits. Relationships between modules and traits are color-coded from red for strong positive correlations to blue for strong negative correlations. For each module, the correlation coefficient and the associated p-value are shown. Correlation p-values shown here treat the 48 libraries as independent and are used for exploration module screening only. The significance of the two infection-associated modules (brown, magenta) was confirmed accounting for oyster identity. Module names appear between the two panels; the bold, colored names correspond to the immune-related modules identified in this study (brown and magenta). B. Mean eigengene expression of the brown (top) and magenta (bottom) modules across infection conditions for each tissue. Points represent mean eigengene values with error bars indicating standard deviation.

